# Neural precursor cells contribute to decision-making by tuning striatal connectivity via secretion of IGFBPL-1

**DOI:** 10.1101/2020.12.29.424678

**Authors:** Erica Butti, Stefano Cattaneo, Marco Bacigaluppi, Marco Cambiaghi, Giulia Scotti, Elena Brambilla, Giacomo Sferruzza, Maddalena Ripamonti, Fabio Simeoni, Laura Cacciaguerra, Aurora Zanghì, Angelo Quattrini, Riccardo Fesce, Paola Panina-Bordignon, Francesca Giannese, Davide Cittaro, Tanja Kuhlmann, Patrizia D’Adamo, Maria Assunta Rocca, Stefano Taverna, Gianvito Martino

**Affiliations:** Neuroimmunology Unit, Division of Neuroscience, Institute of Experimental Neurology (INSPE), IRCCS San Raffaele Hospital, Milan, Italy; Department of Neurosciences, Biomedicine and Movement Sciences, University of Verona, Verona, Italy; Omics Sciences Center, IRCCS San Raffaele Hospital, Milan, Italy; Neuroimaging Research Unit, Institute of Experimental Neurology (INSPE), Division of Neuroscience, IRCCS San Raffaele Hospital, Milan, Italy; Neuropathology Unit, Institute of Experimental Neurology (INSPE), Division of Neuroscience, IRCCS San Raffaele Hospital, Milan, Italy; Department of Biomedical Sciences, Humanitas University, Pieve Emanuele, Milan, Italy; Institute of Neuropathology, University Hospital Muster, Muster, Germany; Molecular genetics of Intellectual Disability, Division of Neuroscience, IRCCS San Raffaele Hospital, Milan, Italy; Vita-Salute San Raffaele University, Milan, Italy

## Abstract

The adult brain retains endogenous neural stem/precursors cells (eNPCs) in two major neurogenic niches, the subgranular zone (SGZ) in the hippocampus and the subventricular zone (SVZ). While in humans and rodents the SGZ contributes to memory functions by generating new neurons that integrate into hippocampal circuitry, the role of eNPCs of the SVZ is less clear. SVZ-eNPCs contribute to the generation of new neurons fated to the olfactory bulb (OB) in rodents but not in humans where they are thought to contribute to striatal neurogenesis as a result of tissue damage.

Here, we show in mice and humans a novel non-neurogenic physiological role of adult SVZ-eNPCs in supporting cognitive functions by regulating striatal neuronal activity. We first provide evidence that GABAergic transmission between parvalbumin-expressing fast-spiking interneurons (FSIs) and medium spiny neurons (MSNs) is tuned by SVZ-eNPCs via secretion of Insulin-Like Growth Factor Binding Protein Like 1 (IGFBPL-1) that, in turn, regulates the Insulin-Like Growth Factor (IGF-1) signalling cascade. Consistently, selective ablation of SVZ-eNPCs and in vivo disruption of the IGF-signalling determine the impairment of intrastriatal coherence. A finding associated with a higher failure rate of GABAergic transmission mediated by FSIs and with striatum-related behavioural dysfunctions impairing decision making. Human validation studies revealed IGFBPL-1 expression in the SVZ and in foetal and induced-pluripotent stem cell-derived NPCs as well as a strong correlation in neurological patients between SVZ pathological damage, reduction of striatum volume and impairment of information processing speed.

All in all, our results highlight a novel non-neurogenic homeostatic role exerted by SVZ-eNPCs on striatal GABAergic neurons that might contribute to cognitive processes involving decisions-making tasks.

## Introduction

Endogenous neural stem/precursor cells (eNPCs) are present in two specific neurogenic niches of the brain, the subgranular zone (SGZ) in the dentate gyrus (DG) of the hippocampus and the subventricular zone (SVZ) adjacent to the lateral ventricles ^1 2 3^. While neurogenesis occurring in the SGZ during adult life is considered to be a crucial physiological process aimed at maintaining memory circuits integrity and cognitive adaptability, in both humans and rodents, less clear is the physiological role of SVZ-eNPCs ^4^. While in rodent SVZ-eNPCs account for olfactory bulb (OB) granule cell renewal, the integration of neurons in the adult human OB accounts for less than 1% of the total neurons exchanged over a century ^5^. At the same time, there is evidence that SVZ-eNPCs might account in humans for striatal neurogenesis, as a reactive process to tissue damage ^6^. Considering these differences, a possible explanation can rely on evolutionary changes in volume and functional performance of the different brain areas in humans compared to rodents but can be also explained by the fact that SVZ-eNPCs, that persist in an undifferentiated state, might exert also non neurogenic functions. Those functions, namely the possibility to promote neuroprotection via the release of soluble molecules ^7 8 9 10 11^, have been recently shown to be exerted by SVZ-eNPCs in pathological conditions but little is yet known about the possibility that such cells might also promote non neurogenic functions in the adult brain in physiological conditions.

The potential contribution of SVZ-eNPCSs to striatal functioning is supported by several evidence. From an anatomical perspective, SVZ-eNPCs are not only in close contact with blood and cerebrospinal fluid (CSF) compartments ^12^ but are also located very closely to and interconnect with the striatum, the main input area of the basal ganglia, whose relevance in regulating cognitive processes, and in particular those involved in decision-making, is increasingly appreciated. Indeed, the ventral striatum primarily mediates motivational processes such as reward and hedonia, whereas the dorsal striatum primarily mediates learn and discrimination ability as well as and engagement with environmental stimuli ^13^ leading to rewards ^14^. Actually, the reward circuit is partially overlapping in both ventral and dorsal striatum ^15^.

This regulatory activity is possible because the striatum is very much interconnected, functionally and anatomically, with the prefrontal cortex, receives massive excitatory input from cortex and thalamus and contains an intricate network of GABAergic synaptic connections projected by several cells types such as medium spiny neurons (MSNs) and local interneurons ^16 17 18^. Among local interneurons, fast-spiking PV-interneurons (FSI) play a key role since they form perisomatic synapses with MSNs, thus resulting in a strong inhibitory activity ^19^.

All in all, those findings prompted us to hypothesize that SVZ-eNPCs might exert a non-neurogenic physiological role on striatal areas involved in cognitive processes. We found that ablation of SVZ-eNPCs in mice caused anatomical changes in striatal MSNs and functional deficits characterized by an altered GABAergic synaptic transmission specifically mediated by FSIs onto MSNs. A finding paralleled by less efficient striatal-dependent cognitive tasks – i.e. as discrimination, learning and reward – involved in decision making ^20 21^. These effects were induced by SVZ-eNPC-mediated decreased secretion of insulin-like growth factor-binding protein-like 1 (IGFBPL-1), a protein that binds and stabilizes insulin-like growth factor 1 (IGF-1) ^22 23^. Finally, we demonstrated that human foetal, adult and induced pluripotent stem cell (iPS)-derived NPCs do significantly express and secrete IGFBPL-1 and showed that pathological injury of the SVZ in humans with neurodegenerative disorders significantly correlates with striatal dysfunction and cognitive impairment affecting decision-related domains. Altogether, these data suggest a new mechanism used by SVZ-eNPCs to exert important non-neurogenic homeostatic ‘cognitive’ functions in striatal areas involved in decision-making.

## Results

### The absence of SVZ-eNPCs causes striatal morphological alterations

In order to investigate the effects of eNPC ablation in the SVZ on nearby striatal neurons, we used a transgenic mouse model expressing thymidine kinase (TK) under the second intron enhancer of Nestin. In these mice (i.e., NestinTK^+^), the SVZ-eNPCs can be selectively ablated upon subcutaneous ganciclovir (GCV) administration ^24^. After GCV administration for 4 weeks in NestinTK^+^ mice (i.e., NestinTK^+^GCV^+^) we observed a strong reduction of SVZ neuroblasts (DCX^+^ cells) and transit amplifying cells (BrdU^+^ cells) compared to controls, namely NestinTK^-^GCV^+^ mice (Fig. 1a, b, c). We next investigated morphological alterations in striatal neurons after eNPC ablation. A morphometric Sholl analysis performed on Golgi-stained MSNs in the striatum of NestinTK^+^GCV^+^ mice revealed a significant increase in the number of dendritic intersections that was paralleled by an increased total length of dendrites (Fig. 1d, e, f) compared to control mice ^25,26^. Moreover, the average spine length was increased in NestinTK^+^GCV^+^ mice (Fig. 1g), while the spine density remained unchanged (Fig. 1h). In addition, using EM analysis we observed that most of the neuronal spines of striatal neurons from NestinTK^+^GCV^+^ mice displayed an elongated, filopodia-like, morphology compared to spines from control mice (Fig. 1i).

**Fig. 1.**
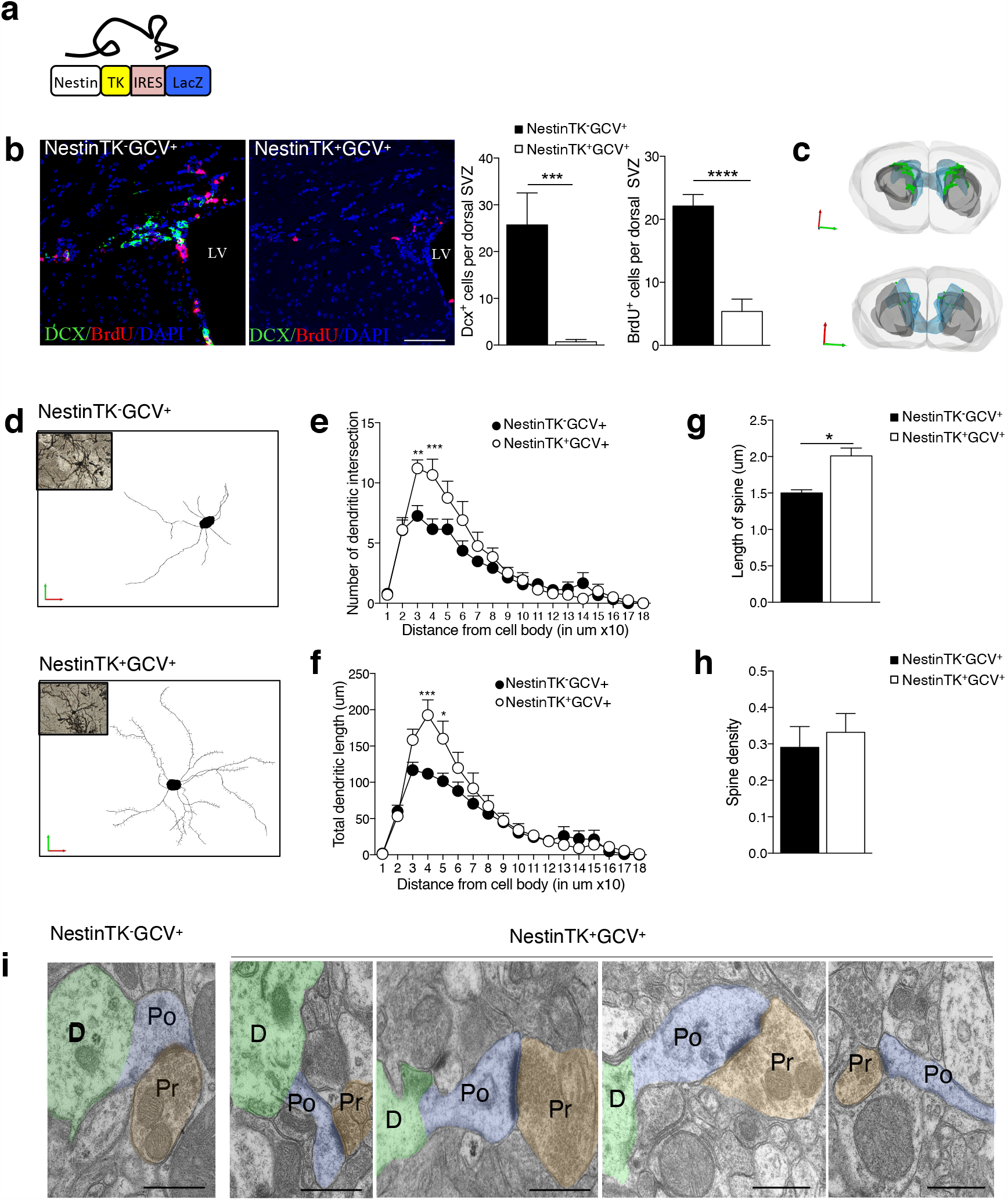
SVZ-eNPC ablation induces structural modifications of striatal MSNs. **a**, Schematic representation of transgenic mouse model with structure of gene expression cassette. **b**, Representative images and quantification of eNPC ablation after 4 weeks of GCV treatment in NestinTK^-^GCV^+^ and NestinTK^+^GCV^+^ mice: neuroblasts were labelled by doublecortin (DCX, in green) and transient amplifying cells for BrdU (in red). Nuclei in blue were counterstained by DAPI. LV, lateral ventricle. Scale bar: 50 μm. n= 3 mice per group. Values represent means ± SEM. ***p=0.0006; ****p=8.9*10^−5^. unpaired two-tailed t-test. **c**. Representative 3D reconstructions of the forebrain staining for DCX (cells in green) of a representative NestinTK^-^ GCV^+^ (up) and NestinTK^+^GCV^+^ (down) mouse after 4 weeks of GCV treatment. Brain hemispheres in light grey, ventricles in blue and striata in grey. **d**. Renderings obtained on Neurolucida of reconstructed MSNs from a representative NestinTK^-^GCV^+^ (up) and NestinTK^+^GCV^+^ (down) mouse. **e, f**, Sholl analysis comparing tracings of medium spiny neurons (MSNs) from NestinTK^-^GCV^+^ and NestinTK^+^GCV^+^ n=6-8 neurons per mice for a total of 3 mice per group. Values represent means ± SEM. In d **p=0.0055, ***p=0.0008, in e *p=0.0144, ***p=0.0001 two-way-ANOVA test followed by Sidak post-test. **g, h**, Striatal MSNs spine length **(g)** and spine density **(h)** in NestinTK^-^GCV^+^ compared to NestinTK^+^GCV^+^ mice. Values represent means ± SEM. In **g** *p=0.0016 Unpaired two-tailed t-test. **i**, Spine morphology analysed by electron microscopy of a representative NestinTK^-^GCV^+^ and a NestinTK^+^GCV^+^ mouse. Colours shown are pseudo colours: dendrites (green), spine (blue) and the synaptic boutons (brown). Scale bar: 0.5 μm.

Increased spine length and augmented short dendritic ramifications of MSNs from NestinTK^+^GCV^+^ mice strongly indicate that the ablation of eNPCs in the SVZ affects neuronal structure. As an additional confirmation of these results, neuronal modifications were reverted back to normality when NestinTK^+^GCV^+^ were allowed to recover by interrupting GCV administration for 4 weeks (i.e., recovery phase) (Extended Data Fig.1). Neither at the end of ablation nor after the recovery phase we observed caspase-3 positive cells or any inflammatory reaction to the GCV procedure in the striatum (data not shown) thus excluding a possible biasing effect on neuronal loss in the striatum.

### SVZ-eNPCs ablation impairs discriminative delay-conditioning task

We next investigated whether the structural and functional alterations of MSN connections observed in NestinTK^+^GCV^+^ mice were paralleled by cognitive impairment sustained by striatum-dependent behaviours. We thus adopted a discriminative delay-conditioning task that is able to assess decision-making tasks in which striatal areas play a major supporting role such as learning to detect, discrimination and reward ^27^. Indeed, mice have to learn that a food pellet is delivered only in association with a tone A (food-reinforced conditioned stimulus: CSA^+^) and not with a non-reinforced tone B (CSB^−^) and must learn, over trials and days, to discern the two stimuli and react appropriately (Fig. 2a). In particular, the appropriate response consists in poking their nose at the food delivery spot at a fixed time delay (20 sec) after the start of the CS.

**Fig. 2.**
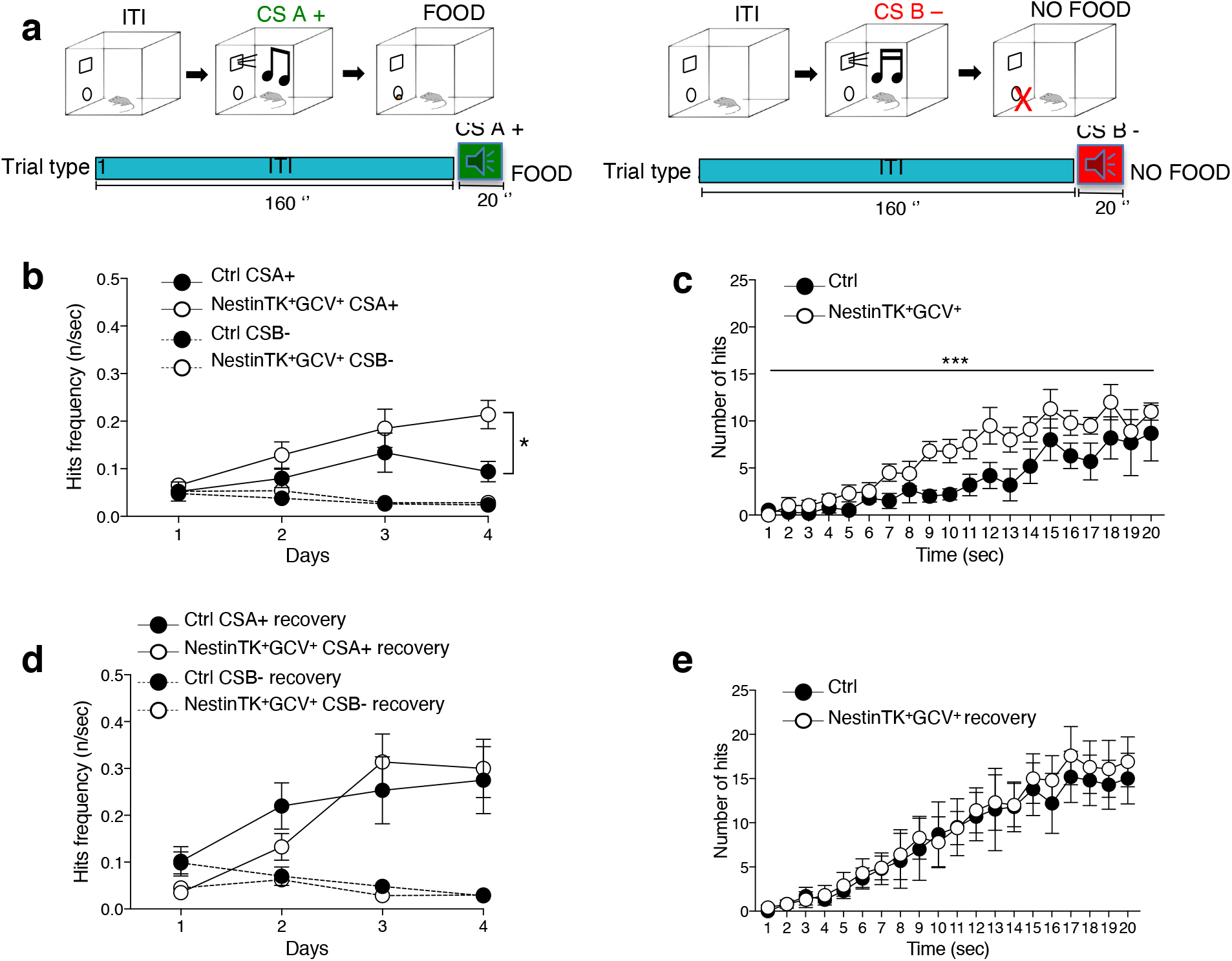
SVZ-eNPC ablation alters time learning in NestinTK^+^GCV^+^ mice. **a**, Schematic representation of the Discriminative Delay-Conditioning protocol used. **b, c**, The graphs show the average frequency of hits during the 20-s CS (A+ or B-) tone **(b)** and the distribution of the hits in successive 1-s intervals during the CS-A+ tone **(c)** in Ctrl and NestinTK^+^GCV^+^ at the end of GCV administration. Panels **d** and **e** display the average frequency **(d)** and the distribution of hits during the CS **(e)** in Ctrl and NestinTK^+^GCV^+^ one month after the end of GCV administration (wash out period). n=12 mice per group. Values represent the mean ± SEM. One way ANOVA genotype effect: F[1,11]=8.3; *p=0.01 in **b**. Kolmogorov-Smirnov test for two data sets ***p<0.0001 in **c**. No significant differences in **d** or **e**.

In this procedure, both NestinTK^+^GCV^+^ and control mice (i.e., NestinTK^-^PBS^+^, NestinTK^-^GCV^+^ and NestinTK^+^PBS^+^) acquire discriminative response to the CSA^+^ over CSB^-^ (Fig. 2b). However, a significant difference between groups appeared clearly by day 4: in response to the CS^+^ that predicts a biological relevant stimulus (food pellet delivery), the number of pokes was greater in NestinTK^+^GCV^+^ mice compared to controls (Fig. 2b), as they repetitively poke their nose well before the appropriate delay had lapsed (Fig. 2c). The analysis of hits distribution during the 20 second CSA^+^ presentation revealed a significant increase in the number of hits in NestinTK^+^GCV^+^ and an overall anticipated (inappropriate) action.

Thus, in a discriminative delay-conditioning task, a behaviourally complex goal tracking procedure, NestinTK^+^GCV^+^ mice are less efficient in engaging and correctly delaying the appropriate response (Fig. 2c). The same procedure was performed in an independent cohort of NestinTK^+^GCV^+^ mice, after one month of recovery, and no difference was found between NestinTK^+^GCV^+^ and control mice (Fig. 2d, e).

### eNPC ablation impairs GABAergic transmission in striatal MSNs

To further investigate whether such striatal alteration had a neurophysiological equivalent, we recorded the oscillatory activity of local field potentials (LFPs) in both striata to measure striatum-striatum coherence with respect to somato-sensorial cortices within the same animals. We observed an increased interhemispheric coherence from 2 to 20 Hz in the striatum of NestinTK^+^GCV^+^ but not in the somato-sensory cortices (Extended Data Fig. 2).

Then, to test whether eNPC ablation affects GABAergic synaptic transmission in striatal slices, we performed whole-cell patch clamp recordings in MSNs to detect spontaneous inhibitory postsynaptic currents (sIPSCs) in the presence of the AMPA-receptor antagonist NBQX (5μM) (Fig. 3a). Average frequency and amplitude of sIPSCs were significantly reduced in striatal MSNs of NestinTK^+^GCV^+^ mice as compared to controls (Fig. 3b, c). In slices prepared from recovered NestinTK^+^GCV^+^ mice (after a 2-months period of recovery after GCV administration), the frequency of sIPSCs had returned to control values, while amplitude remained reduced. Thus, eNPC ablation appeared to induce a partially reversible defect in striatal GABAergic transmission targeting MSNs.

**Fig. 3.**
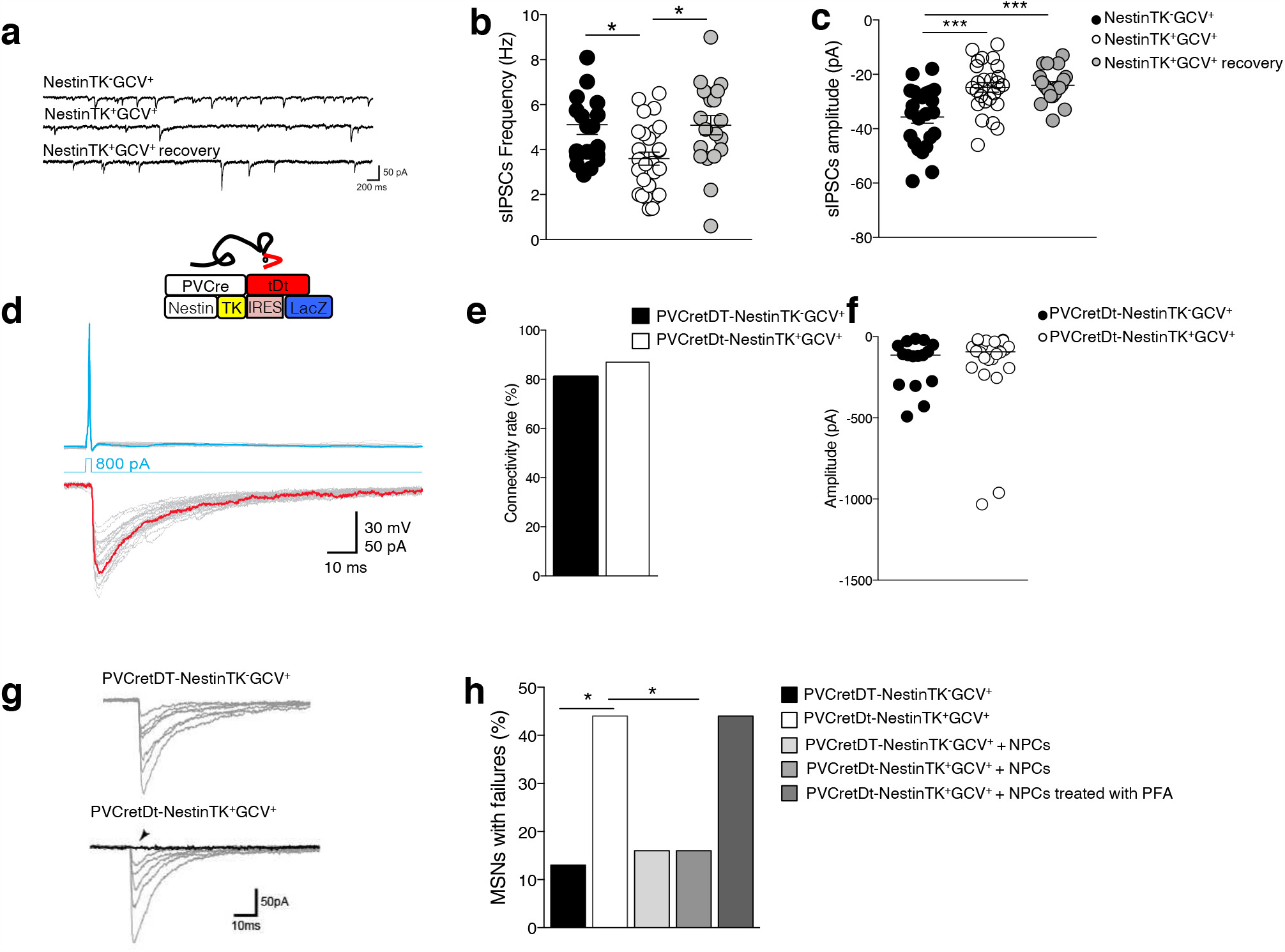
SVZ-eNPC ablation induces deficits in striatal GABAergic synaptic transmission. **a**, Examples of sIPSC recordings in striatal MSNs in slices from NestinTK^-^GCV^+^ mice (top), NestinTK^+^GCV^+^ mice (middle), and NestinTK^+^GCV^+^ mice after a wash out period of two months (bottom). **b, c**, Summary plots showing sIPSC frequencies **(b)** and peak amplitudes **(c)** for the three mice groups. Values represent means ± SEM. *p=0.015 in **b** and ***p=0.0002 in **c**; one-way ANOVA followed by Tukey post-test. n=19 cells in 5 NestinTK^-^GCV^+^, n=27 cells NestinTK^+^GCV^+^ in 5 mice and n=20 cells in 2 NestinTK^+^GCV^+^ mice after a wash out period. **d**, Schematic representation of transgenic mouse model with structure of gene expression cassette. Examples of dual patch clamp recordings showing unitary IPSCs (bottom traces) evoked in an MSN in response to single action potentials (top traces) elicited by injection of a brief suprathreshold current step (800pA, 2ms) in a synaptically connected FSI. The inset shows a schematic cartoon representing a presynaptic FSI and a postsynaptic MSN simultaneously recorded. **e**, Summary histogram of connectivity rates (i.e., percentage of success in finding synaptically connected cells) for FSI-MSN pairs in PVCretDt-NestinTK^-^GCV^+^ (n=8 mice) and PVCretDt-NestinTK^+^GCV^+^ mice (n=12 mice) 15 out of 19 (79%), and 25 out of 30 (83%) pairs respectively. **f**, Summary plot of peak amplitude for FSI-MSN pairs in PVCretDt-NestinTK^-^GCV^+^ and PVCretDt-NestinTK^+^GCV^+^ mice. Values represent medians. Mann-Whitney U test. n=15 and n=25 pairs in PVCretDt-NestinTK^-^GCV^+^ and PVCretDt-NestinTK^+^GCV^+^ mice, respectively. **g**, Examples of unitary IPSCs recorded in a MSN in response to individual APs evoked in a FSI (not shown). Note the presence of failures (arrowhead) in the MSN from an eNPC-ablated mouse. **h**, Summary histogram with percentages of MSNs showing at least one failure in response to individual APs elicited in the connected FSI. Z-Test for 2 population proportions *p=0.045 PVCretDt-NestinTK^-^GCV^+^ vs PVCretDt-NestinTK^+^GCV^+^ and *p=0.046 PVCretDt-NestinTK^+^GCV^+^ vs PVCretDt-NestinTK^+^GCV^+^ NPCs. In the histogram, from left to right, the number of MSNs showing at least one failure are: 2 out of 15 (13%, n=8 mice), 11 out of 25 (44%, n=12 mice), 1 out of 6 (17%, n=2 mice), 3 out of 19 (16%, n=9 mice) and, 4 out of 9 (44%, n=3 mice).

Since sIPSCs may result from GABA release from several unidentified neighbour cell types (interneurons or other MSNs), we performed simultaneous dual whole-cell recordings in identified, synaptically connected pairs of striatal cells. Since parvalbumin-expressing fast-spiking interneurons (FSIs) represent one of the principal sources of local GABAergic projections to MSNs ^28^, we first investigated the properties of FSI-MSN pairs using dual recordings. To visually identify striatal FSIs, we crossed NestinTK mice with mice expressing cre-dependent, floxed tdTomato (tdT) red fluorescent protein under the promoter for parvalbumin (PV; Fig. 3d). For each FSI-MSN pair, unitary IPSCs (uIPSCs) were recorded in the MSN in response to individual action potentials (APs) elicited in the FSI. We found no changes in connectivity rates for FSI-MSN pairs from control vs. PVcre-tdT-NestinTK^+^GCV^+^ mice (Fig. 3e). Similarly, average peak amplitudes of uIPSCs were not significantly different after eNPC ablation (Fig. 3f). However, we found a significantly larger percentage of FSI-MSN pairs showing transmission failures in slices from PVcre-tdT-NestinTK^+^GCV^+^ mice as compared to controls (Fig. 3g, h). Only 13% (2 out of 15) of control FSI-MSN pairs displayed at least one failure in response to a series of 30 individual APs elicited in the presynaptic FSI. This rate was significantly larger (44%; 11 out of 25) in NPC-ablated slices. Interestingly, relative failure rates were restored to control values after adding NPCs directly onto PVcre-tdT-NestinTK^+^GCV^+^ slices (Fig. 3h). Inactivation of NPCs by pre-treatment with paraformaldehyde disrupted, as expected, their ability to restore normal failure rates (Fig. 3h). Altogether, these results suggest that eNPC ablation reduces the ability of striatal FSIs to release GABA onto MSNs. Such impairment was specific of FSI-mediated GABAergic transmission, since connectivity properties of MSN-MSN and somatostatin-expressing interneurons (SOM)-MSN pairs were unmodified after eNPC ablation (Extended Data Fig. 3 and 4, respectively).

Analysis of IPSC cumulative amplitude in FSI-MSN pairs revealed a significantly smaller size of the readily releasable vesicle pool (RRP) in eNPC-ablated slices than controls (Extended Data Fig. 5). In addition, confocal imaging analysis revealed a small but significant reduction in PV-positive GABAergic synaptic terminals contacting putative MSNs (Extended Data Fig. 5), suggesting that reduced FSI-mediated GABA release is caused by a reduced availability of presynaptic release sites in eNPC-ablated mice.

To further analyse this phenomenon and, although improbable, to avoid biassing effects due to GCV administration, we also analysed aged mice (18 months) that are characterised by a spontaneous progressive reduction of SVZ-eNPCs ^29 30^ (Extended Data Fig. 6). As expected, in these mice we also observed a decreased neurogenesis using both quantitative real time PCR and immunofluorescence staining using NPCs markers (DCX and BrdU; Extended Data Fig. 6a, b). Interestingly, average frequency and amplitude of sIPSCs recorded in MSNs were significantly lower in aged than young mice (Extended Data Fig. 6c). Accordingly, we found a much higher rate of MSNs showing failures in FSI-MSN pairs recorded in the aged group compared to younger animals (Extended Data Fig. 6d) thus reinforcing the concept that eNPC loss is associated with a defect in GABA release from striatal FSIs.

### IGFBPL-1 is down-regulated in NestinTK^+^GCV^+^

In order to unravel possible mechanisms underlying the striatal impairment after eNPC ablation we performed RNA-sequencing (RNA-seq) on SVZ. A differential gene expression (DGE) analysis was accomplished to compare NestinTK^+^GCV^+^ with controls mice: we identified 2 sets of unique genes that were differentially down-regulated (n=115) or up-regulated (n=66) in NestinTK^+^GCV^+^ samples, setting a significance threshold of P ≤ 0.01 and a difference greater than 2 fold (Fig. 4a, b, c). Gene set enrichment analysis (GSEA) identified a strong enrichment in genes involved in the matrisome, with *Igfbpl-1 −* a binding protein that stabilizes the Insulin-like Growth Factor-1 (IGF-1) and allows it to bind its receptor - as one of the top down-regulated genes (LogFC= −5,97) (Fig. 4d). Strong down-regulation of *Igfbpl-1* was further validated by rt-PCR in NestinTK^+^GCV^+^ mice (Fig. 4e), and in aged mice compared to young mice (Extended Data Fig. 6e).

**Fig. 4.**
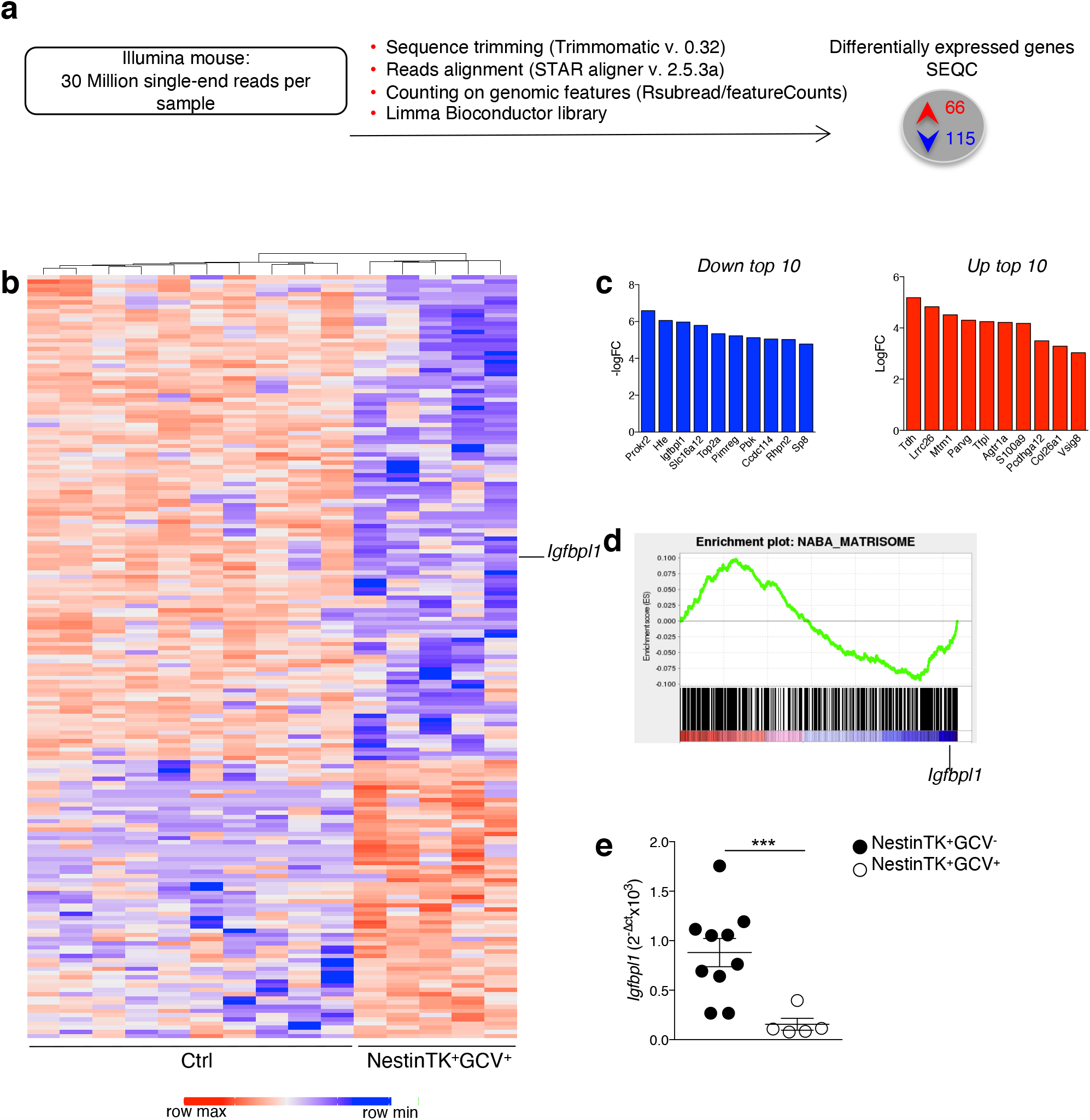
Gene expression analysis in the SVZ in NestinTK^+^GCV^+^ mice. **a**, Workflow of RNA-seq gene expression analysis in control and NestinTK^+^GCV^+^ treated mice. **b**, Heatmap with hierchical clustering of expression values of 115 down regulated genes and 66 up regulated genes with ≥2-fold change in control and NestinTK^+^GCV^+^ mice. (blue means downregulation, red upregulation) n=5 mice per group. **c**, The graphs represent the top 10 downregulated (in blue) and upregulated (in red) genes. **d**, GSEA curve of genes expressed differentially in NestinTK^+^GCV^+^ mice compared to control mice. Enrichment plot for NABA-Matrisome. (FDR= 0.04; NES= −2.5). **e**, Quantitative PCR validation of *Igfbpl-1* expressed gene in control mice compared with NestinTK^+^GCV^+^ mice. n=10 ctrl mice and n=5 NestinTK^+^GCV^+^ mice. Values represent means ±SEM. ***p=0.0027; Mann-Whitney U test.

We then looked for IGFBPL-1 distribution within the adult brain using immunofluorescence imaging which revealed that eNPCs display a strong expression for IGFBPL-1 in the SVZ, along the lateral ventricle, the rostral migratory stream up to the olfactory bulbs, while it was slightly detectable in the striatum. No evidence of IGFBPL-1 was however found in the cortex and in the thalamus (Fig. 5a). eNPC ablated mice showed lack of IGFBPL-1 thus confirming at protein level the data obtained on gene expression level (Fig. 5b).

**Fig. 5.**
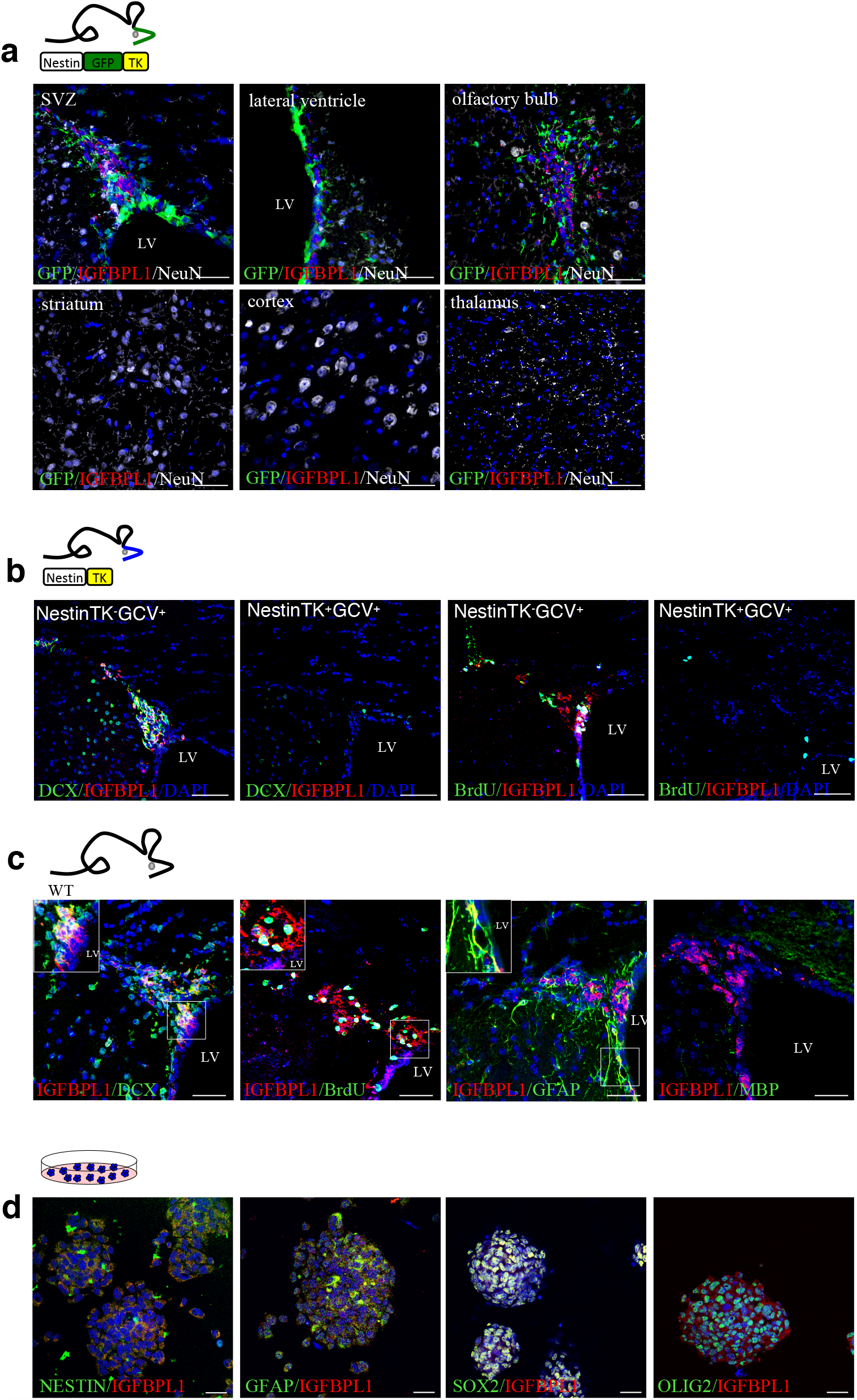
IGFBPL-1 expression in the mouse SVZ. **a**, Representative confocal images of NestinGFPTK coronal brain sections obtained at the level of the dorsal SVZ, of the ventral SVZ (lateral ventricle), of the rostral migratory stream at the level of the olfactory bulbs, of the striatum, cortex and thalamus. Slides were labelled for GFP in green, IGFBPL-1 in red and NeuN in white. Nuclei in blue were stained by DAPI. Scale bar: 50 μm. **b**, Representative confocal images of the SVZ of NestinTK mice either treated or not with GCV and stained for DCX or BrdU in green, IGFBPL-1 in red. Nuclei in blue were stained by DAPI. Scale bar: 50 μm. **c**, Representative confocal images of the SVZ of wt mice labelled for IGFBPL-1 in green, DCX, BrdU, GFAP and MBP in red. Nuclei in blue were stained by DAPI. The inserts (white square) represent the double positive cells. Scale bar: 50 μm. **d**, Representative confocal images of *in vitro* neurospheres labelled for IGFBPL-1 in red, NESTIN, GFAP, SOX2 and OLIG2 in green. Nuclei in blue were stained by DAPI. Scale bar: 25 μm.

In order to understand which neural-derived cell type of the SVZ produces and releases this binding protein, we analysed subtypes of eNPCs such as neuroblasts (DCX^+^), type-C cells (BrdU^+^), type-B cells (GFAP) and oligodendrocytes (MBP). We found that all subtypes of eNPCs, but not oligodendrocytes, express the protein that is visible in the extracellular space close to the cell body (Fig. 5c). Moreover, we observed that mouse-derived neurospheres *in vitro* express IGFBPL-1 protein, as expected (Fig. 5d).

### IGFBPL-1 and IGF pathways are impaired in the striatum of eNPC-ablated mice

In order to assess the downstream effects of reduced expression of eNPC-secreted IGFBPL-1, we analysed the expression profile of the striatum in NestinTK^+^GCV^+^ vs. control mice.

A list of 184 differentially expressed genes with a P ≤ 0.01 and a difference greater than 2-fold at the same time (Extended Data Fig. 7a, b) was generated and, according to Pre-ranked GSEA, a significant enrichment was observed for the *Igf-1* Reactome pathway with genes downregulated in NestinTK^+^GCV^+^ compared to control mice (Extended Data Fig. 7c).

Moreover, both in situ hybridization and rt-PCR revealed abundant expression of *Igf-1r* in the striatum that was increased in NestinTK^+^GCV^+^ compared to control mice (Extended Data Fig. 7d, e and paralleled by decreased IGF-1 protein level in the SVZ (Extended Data Fig. 7f).

### A key role of IGFBPL-1 in maintaining the striatal equilibrium

In order to verify the role of IGFBPL-1 we generated a NPCs line infected with a lentivirus expressing a short interfering RNA for *Igfbpl-*1 (NPCs-shIgfbpl-1) (Fig. 6a). We then applied NPCs-shIgfbpl-1 on striatal slices obtained from NPC-ablated NestinTK^+^GCV^+^ mice. The average sIPSC frequency recorded in MSNs after addition of NPCs-shIgfbpl-1 was significantly lower than control values, while failure rates in FSI-MSN pairs were significantly higher than controls and similar to ablated mice slices (Fig. 6b, c, d; cf. Fig. 4h). Moreover, the average sIPSC frequency recorded in MSNs of NestinTK^+^GCV^+^ mice was significantly increased after incubation with IGF-1 (10nM; Fig. 6c). Accordingly, the rate of MSNs showing failures in FSI-MSN pairs recorded after addition of IGF-1 was similar to controls (Fig. 6d).

To confirm the crucial role of IGFBPL-1 in the striatal functionality, we used a transgenic *Igfbpl-1*^*-/-* 22^ mouse line in which the neurogenesis in the SVZ is maintained, but the eNPCs do not expressed IGFBPL-1 (Fig. 6e). *Igfbpl-1*^*-/-*^ mimicked what we observed in NestinTK^+^GCV^+^ as they showed a significant increase in the number of dendritic intersections and of the total length of dendrites compared to wild-type mice (Fig. 6f, g). Spine length and spines density remained unchanged (Fig. 6h, i). The average sIPSC frequency recorded in MSNs was significantly lower in *Igfbpl-1*^*-/-*^ mice than control mice, while the amplitude did not change (Fig. 6j, k). These data suggest that NPCs regulate the function of FSI-MSNs synapses through IGFBPL-1 release.

**Fig. 6.**
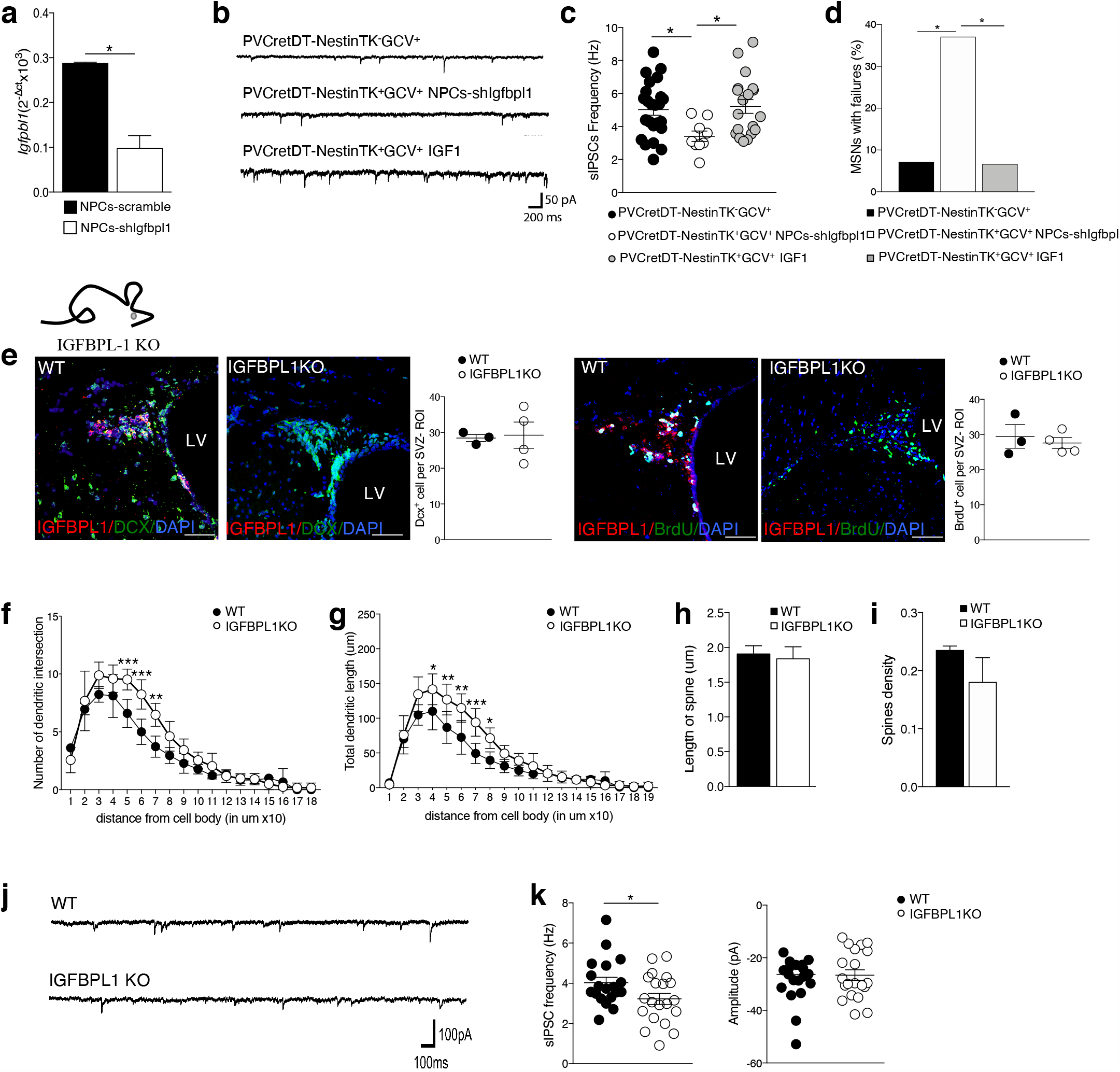
*Igfbpl-1* loss of function in NPCs and *Igfbpl-1*^*-/-*^ mice recapitulate morphological and neurophysiological deficits caused by SVZ-eNPC ablation. **a**, The graph shows a quantification of the *Igfbpl-1* in NPCs infected with a lentivirus expressed sh-*Igfbpl-1*. Values represent means ± SEM *p=0.011, unpaired two-tailed t-test. **b**, Examples of sIPSCs recorded in striatal MSNs in NPC-ablated slices before (top) and after IGF-1 incubation (10μM). **c**, Summary dot plots with average sIPSC frequencies. Values represent the mean ± SEM. *p=0.023 (PVCretDt-NestinTK^+^GCV^+^ vs PVCretDt-NestinTK^+^GCV^+^NPCs-shIgfbp-1, *p=0.038 (PVCretDt-NestinTK^+^GCV^+^NPCs-shIgfbpl-1 vs PVCretDt-NestinTK^+^GCV^+^ IGF1); one-way ANOVA followed by Tukey post-test. n=5 PVCretDt-NestinTK^-^GCV^+^ mice (n=24 cells), n=6 PVCretDt-NestinTK^+^GCV^+^ mice for NPCs-shIgfbpl-1 application (n= 9 cells), n=3 PVCretDt-NestinTK^+^GCV^+^ mice with IGF-1 incubation (n=19 cells). **d**, Summary histograms comparing reliability of FSI-MSN GABAergic synapses (measured as % of MSNs displaying at least one failure after FSI activation). Z-Test for 2 population proportions; *p=0.04 (PVCretDt-NestinTK^+^GCV^+^ vs PVCretDt-NestinTK^+^GCV^+^ NPCs-shIgfbpl-1), *p=0.032 (PVCretDt-NestinTK^+^GCV^+^ vs PVCretDt-NestinTK^+^GCV^+^ IGF1). From left to right, the number of MSNs showing at least one failure was: 1 out of 14 (7%, n= 5 mice), 10 out of 27 (37%, n= 6 mice) and 1 out of 15 (7%, n= 3 mice). **e**, Representative images and quantification of neurogenesis in WT C57Bl/6 and *Igfbpl-1-/-* mice: neuroblasts were labelled for doublecortin (DCX, in green on the left) and transient amplifying cells for BrdU (in green on the right) while IGFBPL-1 (in red). Nuclei in blue were counterstained by DAPI. The lateral ventricle is denoted as LV. Values represent means ± SEM. n= 4 mice per group. Scale bar: 50 μm. **f, g**. Sholl analysis comparing tracings of medium spiny neurons (MSNs) from C57Bl/6 and *Igfbpl-1-/-* mice. Data are presented as number of dendritic intersections **(f)**, and total dendritic length plotted at increasing distance from the cell body **(g)**. Values represent means ± SEM. In **b** ***p=0.0008, ***p=0.0001, **p=0.002; in **c** *p=0.0473, **p=0.0029, **p=0.0014, ***p=0.0005, *p=0.0465 Two-way-ANOVA test followed by Sidak post-test. n=6-8 neurons per mice for a total of 3 mice per group. **h, i**, Striatal MSNs spine length **(h)**, and spine density **(i)** in C57Bl/6 compared to *Igfbpl-1*^*-/-*^ mice. Values represent means ± SEM. n= 3 mice per group. **j**, Spontaneous IPSCs recorded in WT and *Igfbpl-1*^*-/-*^ mice. **k**, Summary data with average IPSC frequencies and amplitude measured in WT C57Bl/6 and *Igfbpl-1*^*-/-*^ mice. Values represent means ± SEM. *p=0.0435, unpaired two tailed t-test. n=3 mice for group.

### IGFBPL-1 is physiologically expressed in the human brain

To ascertain the expression by human NPC specificity of *Igfbpl1* expression, we quantified the activation of *Igfbpl-1* downloading scATAC-seq data for three cell populations (fibroblasts, iPSs, iPS-NPCs) derived from the two same individuals (Accession ID: E-MTAB-9649, manuscript submitted). We calculated gene activity scores, defined as the sum of the scATAC-seq signal over the gene body extended 2kb upstream the TSS <10.1101/2020.11.09.373613>, for *Igfbpl1, Nestin* and *Pax6*. We found that activity of *Igfbpl-1* is specific to iPS-NPCs (Fig. 7a, upper panel). We also analysed the pseudobulk ATAC profiles over *Igfbpl-1* in the three cell populations, which reveal increased accessibility in regulatory elements in the first intron of *Igfbpl-1*. By looking at the annotation of DNAseI Hypersensitive Sites catalog ^31^ we found that the enrichment happens over regulatory elements annotated for the “Primitive/embryonic” and the “Neural” components (Fig. 7a, lower panel). In accordance with these data, we found that iPS-NPCs do express consistently *Nestin, Pax6* and *Igfbpl-1* m-RNA and produce IGFBPL-1 protein (Fig. 7b).

**Fig. 7.**
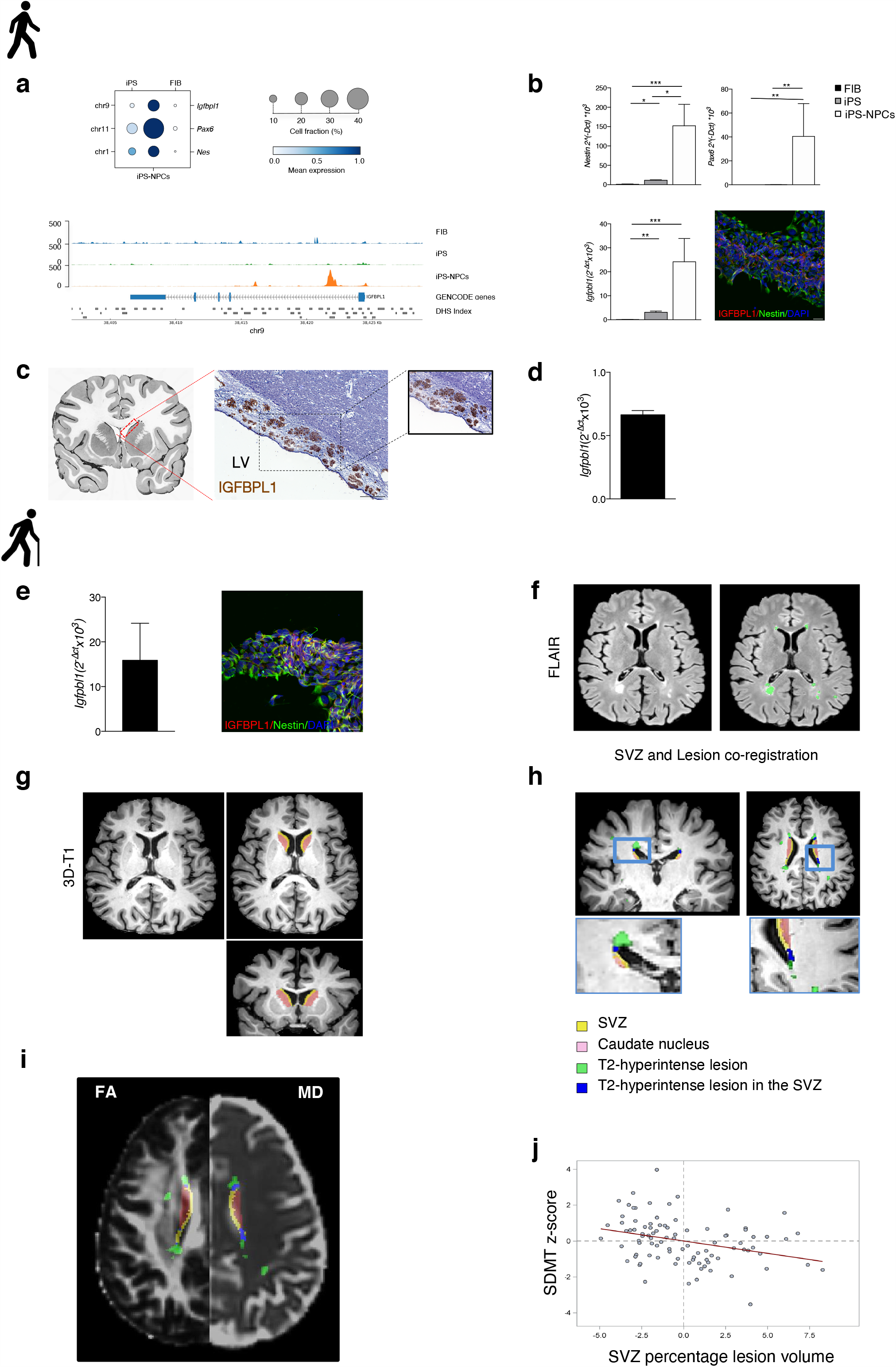
Human SVZ express and secrete IGFBPL-1 and its damage correlates with impaired cognitive performances in patients affected by neurodegeneration. **a**, In the upper panel, dot plot representing the gene activity of selected genes in three cell populations (FIB: fibroblasts, iPS: induced Pluripotent Stem cells, iPS-NPC: Neural Precursor Cells derived from iPS) is shown; each gene is marked by its gene symbol and the genomic coordinates used to derive the gene activity score and the dot size is proportional to the fraction of cells with non-zero signal in the population. Colour code and bubble size represent the mean intensity and % cell fraction of the gene activity score, respectively. In the lower panel, pseudobulk scATAC-seq profiles over *Igfbpl-1* locus in three cell populations; each track represents the average ATAC-seq signal of two individuals in separate cell populations and data values are scaled to the number of cells used to derive the profile. In addition to the *Igfbpl-1* gene structure, the set of DNAseI Hypersensitive Sites (DHS Index) is also represented. **b**, Quantitative PCR of *Nestin, Pax6* and *Igfbpl-1* expressed gene in fibroblasts, iPS and iPS-derived NPCs obtained from three different healthy subjects. Values represent means ± SEM. Immunofluorescent staining showing IGFBPL-1 (in red) and NESTIN (in green) expression in iPS-derived NPCs from healthy subject. Scale bar: 20 μm. **c**, Representative human periventricular brain section showing SVZ cells lining the ventricle consistently expressing IGFBPL-1. Scale bar: 200 μm, 100 μm in the magnification. **d**, Quantitative PCR of *Igfbpl-1* expressed gene in human foetal NPCs. Data shown represent the mean ± SEM of 3 technical replicates. **e**, Quantitative PCR of *Igfbpl-1* expressed gene in iPS-derived NPCs obtained from fibroblasts of three MS patients. Values represent means ± SEM. Immunofluorescent staining showing IGFBPL-1 (in red) and NESTIN (in green) expression in iPS-derived NPCs from MS patient. Scale bar: 20 μm. **f-i**, T2-hyperintense lesion masks were segmented on FLAIR images **(f)** and co-registered on high-resolution T1-weighted sequences **(h)**, fractional anisotropy (FA) and mean diffusivity (MD) maps **(i)**. Caudate nuclei were automatically segmented on high-resolution T1-weighted sequences. According to anatomical references, a subventricular zone (SVZ) mask was segmented on T1-weighted images in the Montreal Neurological Institute space and then registered on native T1-weighted image **(h)**, FA and MD maps **(i)**. Age-, sex- and phenotype-adjusted partial correlation between Symbol Digit Modalities Test (SDMT) and SVZ percentage lesion volume are plotted in **j**. Data from 97 MS patients (mean age 36.8 ± 7.56 years, female/male [F/M] =55/42, median EDSS 2.0, median disease duration 5.0 years) and 43 age- and sex-matched healthy controls (HC, 38.8 ± 6.25 years, F/M=19/24) are reported.

To validate data obtained *in vitro* we also analysed, the expression of IGFBPL-1 in a human brain autoptic tissue obtained from a young patient suffering from a non-central nervous system-confined disorder. IGFBPL-1 was found to be expressed in the ribbon of SVZ lining the lateral ventricles where cells with astrocytic characteristics that can function as NPCs are located (Fig. 7 c). Since the precise location of the stem cells is difficult to establish in the SVZ, to further confirm that *in vivo* NPCs do secrete IGFBLP-1, we finally analyzed human fetal derived NPCs and showed that they do also express *Igfbpl-1* m-RNA (Fig. 7d).

### SVZ damage impairs cognition in patients with MS

We next sought to understand whether in humans, SVZ functional decline is associated to cognitive impairment in the decision-making domain due to striatal dysfunctions. We purposely focussed on patients with MS that have early lesions in particular in the periventricular area with significant tissutal damage. Indeed, the presence of focal lesions abutting lateral ventricles, hence possibly involving the SVZ, is a hallmark of MS. Moreover, cognitive alterations are reported in up to 50% of patients at disease onset ^32^, with information processing speed - an integral and relevant component of the decision-making circuit ^33 34^ - undergoing the earliest impairment ^35^.

We first established that iPS-derived NPCs, obtained from skin fibroblasts from 3 MS patients, expressed IGFBPL-1 at both mRNA and protein level likewise healthy subjects (Fig. 7e, b). We then performed an association study by assessing focal and microstructural damage of the SVZ-eNPC and striatal areas with magnetic resonance imaging (MRI) and cognitive dysfunction with the Symbol Digit Modalities Test (SDMT), in a large cohort of MS patients. SDMT, the most widely used test for assessing information processing speed, was selected as the neuropsychological item most sensitive to cognitive impairment in MS ^36^, and the most administered in MS clinical trials evaluating cognitive disability ^37^. Results were compared to age- and sex-matched healthy volunteers. Ninety-seven MS patients (mean age 36.8 ± 7.56 years, female/male [F/M] =55/42, median EDSS 2.0, median disease duration 5.0 years) and 43 age- and sex-matched healthy controls (HC, 38.8 ± 6.25 years, F/M=19/24) underwent a 3.0 T brain MRI and SDMT testing.

The MRI protocol included fluid attenuation inversion recovery (FLAIR) for T2-hyperintense lesion identification, high-resolution 3D T1-weighted sequence for global and regional brain volumes measurement and diffusion weighted images for the assessment of microstructural integrity (fractional anisotropy and mean diffusivity, correlating with axonal and myelin loss respectively) ^38^.

Starting from anatomical reference ^39^, a mask of the SVZ was obtained from high-resolution T1-weighted images in a standard space (Montreal Neurological Institute space) and then co-registered on native T1- and diffusion-weighted sequences (Fig. 7f-i). The percentage of SVZ volume occupied by lesions was defined as SVZ percentage lesion volume (SVZ percentage LV). In MS, mean SVZ percentage LV was 4.2%. SVZ normal-appearing tissue was characterized by increased mean diffusivity (0.89 vs 0.86, p=0.04) and preserved fractional anisotropy compared to HC.

Although all age-, sex- and phenotype-adjusted partial correlations between caudate volume (normalized for head size) and measures of global brain damage were significant (r absolute values range: 0.34-0.54; p range: <0.0001-0.0008), the stepwise multiple linear regression selected brain volume (p=0.0037), SVZ percentage LV (p=0.0033), mean cortical thickness (p=0.0012) and SVZ intralesional mean diffusivity (p=0.0061) as independent predictors of caudate volume (R^2^=0.58). SDMT performance, rated as education-corrected z-scores ^40^ (median −0.28), correlated only with the regional injury of the SVZ (percentage LV, intralesional fractional anisotropy and normal appearing tissue mean diffusivity, r absolute values range: 0.23-0.31; p range:0.028-0.0028). SVZ percentage LV was selected as the only independent predictor of SDMT performance (p=0.0017, R^2^=0.26), overcoming all other measures of cerebral damage, including brain volume, grey matter volume, white matter volume and cortical thickness.

Altogether our results clearly indicate that SVZ-eNPCs do secrete IGFBPL-1 in both physiological and pathological conditions and that, in patients with MS, SVZ damage is strongly associated with the impairment of information processing speed. This further supports the idea that eNPCs might contributing to cognitive impairment involving decision making in humans.

## Discussion

In rodents, the SVZ has been described mainly as a source of newly formed neurons able to replace granule cells in OB upon migration along the rostral migratory stream ^41^. However, considering the emerging data questioning the existence of a ‘physiological’ neurogenesis in the SVZ (and in the SGZ) of the adult human brain ^42^, the peculiar position of the SVZ itself located several milli(centi)meters far from the OB but very close to basal ganglia, and the evidence supporting non-neurogenic roles for SVZ-eNPCs in both rodents and humans as a result of pathological damage ^43^, it is tempting to speculate that neurogenesis might not be the sole function of SVZ-eNPCs and that non-neurogenic physiological functions could, at least in part, explain this conundrum.

Using dual-patch technology, here we demonstrate that SVZ-eNPCs control GABAergic synaptic connectivity - in particular the unidirectional synapses from FSIs to MSNs - thus representing a crucial player in shaping the morphological structure as well as the functionality of the main neuronal populations residing within the striatal area. Several data support this result. In SVZ-eNPC ablated mice, MSNs showed a significant increase in the number of dendritic intersections, an increase of total length of dendrites and spine and an elongated, filopodia-like, morphology. These alterations were paralleled by a functional derangement of striatal neurons as we observed an increase of glutamatergic currents <23008234> as well as an impairment of GABAergic transmission onto MSNs, due to a decreased ability of FSIs to release GABA. Collectively, our data strongly suggest that the absence of SVZ-eNPCs causes defects in striatal FSIs, a population of cells playing a crucial role in decision-making tasks ^21^ which represents one of the principal sources of local GABAergic projections to MSNs.

Our results are consistent with previous studies showing how striatal PV interneurons regulate striatal-based behavioural tasks ^44 45 46^. The balance between inhibitory and excitatory synaptic transmission is important not only in regulating the mood and feeding behaviour but also to learning, memory and appropriate behavioural responses to external stimuli ^47^. Interestingly, it was recently demonstrated that PV interneurons are also crucial in decision-making tasks, as they influence the initial expression of reward-conditioned responses and their contribution to performance declines with experience ^21^.

The ensuing hypothesis that SVZ-eNPCs are able to control the circuitry of the nearby striatum was further confirmed in a more physiological model of SVZ-eNPC impairment - the aged mouse. It is known that a strong impairment of the neurogenesis occurs in aged mice, probably due to eNPCs cycle dynamic changes ^30^, i.e. the eNPCs acquiring a senescent phenotype ^48^ and shifting to ‘time in quiescence’. In SVZ-eNPC aged mice we observed that the sIPSC frequency recorded in MSNs was significantly lower and the rate of MSNs showing failures in FSI-MSN pairs was higher compared to young mice.

We then explored the molecular mechanism(s) sustaining SVZ-eNPC non neurogenic functions, such as the abovementioned homeostatic control of striatal activity. By means of a gene expression profile study, we found in the striatum of SVZ-eNPC ablated mice a down-regulation of the *Igf-1* pathway. IGF-1 is a growth factor involved in several processes in the brain (i.e. neuronal cell proliferation, survival, axonal growth and synaptogenesis) ^49^ whose paracrine and autocrine mode of action is modulated by IGF-binding proteins ^50^. We better detailed which component of the *Igf-1* pathway was malfunctioning due to the absence of SVZ-eNPCs. Using a RNAseq approach, we found that the *Igfbpl-1* gene was down-regulated in SVZ-eNPCs of ablated mice. To further support this evidence, we showed *in vivo* that all subtypes of eNPCs, but not of mature neuronal cells, express IGFBPL-1 and that SVZ-eNPC ablated mice lack IGFBPL-1 protein expression. Finally, ATAC-Seq analysis confirmed *Igfbpl-1* chromatin-accessibility in a large proportion of iPS-NPCs but not in fibroblasts and iPS further confirming the specificity of our findings. Recent evidence indicating, at single-cell level, that *Igfbpl-1* is expressed in the adult murine V-SVZ by actively dividing NPCs as well as in transit amplifying cells and neuroblasts strongly corroborate our results ^23^.

An active role of IGFBPL-1 in brain functioning is supported by several evidence. Secreted IGFBPL-1 has been shown to be involved in axonal growth via the stabilisation of IGF-1 biding to its receptor ^22^. Transgenic mice overexpressing IGFBPL-1 in brain have impaired brain growth and reduced glial cell proliferation in response to injury ^51^. We thus hypothesized that the lack of such protein might be the culprit of the alterations observed before also considering previous data showing that SVZ-eNPCs have the ability to secrete proteins that, in turn, do exert homeostatic neuroprotective functions within the brain ^52^. To further analyse the effective role in vivo of IGFBPL-1, we first analysed *Igfbpl-1*^*-/-*^ mice. The same morphological and functional alterations observed in the striatum of SVZ-eNPC ablated mice were recorded in *Igfbpl-1*^*-/-*^. Furthermore, patch clamp in vitro studies supported this evidence since *Igfbpl-1* loss of function approaches in NPCs were unable to rescue the IPSCs frequency and the number of FSI-mediated GABA transmission failures back to control values, while the administration of recombinant IGF-1 rescued the electrophysiological deficits. Altogether, these data suggest that IGF-1 - produced within the brain or reaching the brain from the liver trough the bloodstream and choroidal plexus ^53^ - does reach the SVZ where it binds to IGFBPL-1 produced by eNPCs. Once bound to IGFBPL-1, IGF-1 can bind its receptor on the MSNs and FSIs. We can thus suggest that in ablated neurons, which lack IGFBPL-1, IGF-1 availability and binding to its receptor is impaired thus causing the functional deficit in the striatal neurons.

To prove the human relevance of our mouse results, we performed a human validation study. We first tested whether or not human SVZ-eNPC were capable of expressing and secreting IGFBPL-1. We found that autoptic brain material as well as human foetal NPCs and iPS-derived NPCs from healthy controls do express and secrete IGFBPL-1. Furthermore, ATAC studies showed that *Igfbpl-1* is abundantly express by almost 50% of the cells analyzed.

We then measured the correlation between SVZ damage and cognitive impairment in patients with MS. This disease was purposely selected for several reasons. Actually, worse demyelination and tissue loss in periventricular regions ^54-56^, involving the SVZ, are hallmarks of MS. Early cognitive alterations involving the decision making domain - i.e. impairment of information processing speed ^35^ - are reported in up to 50% of MS patients at disease onset ^32^. Unbalanced expression of IGF1 and its receptor – that is found downregulated in MS lesions ^57^ - foster tissue damage in MS patients since they both contribute to myelin production by oligodendrocytes. ^58^ We first confirmed that iPS-derived NPCs from MS patients do express and secrete IGFBPL-1 at comparable level than healthy subjects thus further confirming that the IGF signaling pathway is active in NPCs from MS patients. We then applied advanced MRI sequences to assess microstructural tissue integrity in 96 patients with MS. Results obtained showed that SVZ damage (i.e., SVZ percentage LV, and SVZ intralesional mean diffusivity) is an independent predictor of caudate volume (R2=0.58) and that information processing speed, correlated only with the regional injury of the SVZ. All in all, our data, while confirming mouse results, support the concept that human SVZ-eNPCs might exert (possibly via the IGF signaling pathway) a relevant role in cognitive processes and that such homeostatic role is impaired in patients showing pathological SVZ damage.

In conclusion, here we show for the first time a novel non-neurogenic physiological role of SVZ-eNPCs in supporting GABAergic connectivity within striatal circuits, through the production and the release of IGFBPL-1. This finding suggests that SVZ-eNPCs might contribute, at least in part, to cognitive functions and, in particular, those encompassing decision-related tasks. It is therefore tempting to speculate that SVZ-eNPCs might represent an integral part of the cellular components of cognitive circuits sustaining decision-making processes.

## Methods

### Study approval and animals

Adult female and male C57Bl/6 (6-8 weeks old) aged (18 weeks old) and transgenic mice were purchased from Charles River or generated in our animal facility in SPF conditions. Experimental procedures, performed blindly, were approved by the Institutional Animal Care and Use Committee (no. 750 and 798) of the San Raffaele Scientific Institute, Milan (Italy). To selectively ablate SVZ eNPCs, we used the NestinTK transgenic mouse line, in which the thymidine kinase gene is under the control of the second intron of Nestin rat promoter ^24^. Treatment of NestinTK, PVcretdT-NestinTK and SomChR-NestinTK mice with ganciclovir for 4 weeks (2 weeks for electrophysiological experiments), by means of a subcutaneously implanted osmotic minipump, induced ablation of Nestin-positive eNPCs. To define the areas of the brain expressing the IGFPBL-1 we used NestinGFPTK transgenic mouse line, in which the NPCs are marked by green fluorescence. For the dual patch clamp recordings, we used PVcretdT-NestinTK obtained in our animal facility by crossing NestinTK with PVcre and Rosa26 Tomato. For detection of somatostatin interneurons, we used SomChR-NestinTK obtained in our animal facility by crossing NestinTK with SomChR. The *Igfbpl-1 ko* line was kindly provided by Dong Feng Chen’s laboratory ^22^.

### Tissue pathology

Mice were deeply anesthetized by intraperitoneal injection of Avertin [2,5g of 2,2,2 tribromoethanol (# T48402 Sigma-Aldrich), dissolved in 5ml of 2-metil-2-butanol (Aldrich) and 200ml of ddH_2_O]. The toe pinch-response method was used to determine the depth of anaesthesia. Mice were first trans-cardially perfused with a saline buffer (0.9% NaCl) plus EDTA [500μl of EDTA (Sigma) 0.5M were added to 250 ml of saline buffer] at room temperature (RT) and then with 4% paraformaldehyde (PFA; # 158127 Sigma-Aldrich) in PBS (Phosphate Buffered Saline) 1X to preserve tissue in a life-like state. Brains were removed and post-fixed in 4% PFA overnight at +4 °C. On the following day, brains were washed in PBS 1X three times for 5 minutes and cryopreserved in 30% sucrose (Sigma) in PBS 1X. After 48 hours brains were embedded in Tissue-Tek OCT compound (BioOptica) and frozen in isopentane in a liquid nitrogen bath. Brains were stored at −80 °C until sectioned. Coronal sections of 14 μm of thickness were obtained using a Leica cryostat (Leica CM1850), collected onto Superfrost slide and air dried overnight before storing them at −80 °C until staining.

For immunofluorescence, brain sections were air dried for at least 40 minutes and then rinsed 3 times for 5 minutes with PBS 1X. Non-specific binding sites were blocked by incubation in the blocking solution (PBS 1X containing 0.1% Triton X-100, 10% FBS, 1 mg/ml BSA) for 1 hour at RT. Sections were then incubated overnight at +4 °C with the appropriate primary antibody diluted in blocking solution in a humidity chamber. The following primary antibodies were used: goat anti DCX (Santa Cruz Biotechnology, product no: sc-8066, lot: A0614 (1:100)), rat anti BrdU (Abcam, product no: ab6326, (1:100)), rabbit anti DCX (Abcam, product no: ab77450, (1:100)), goat anti IGFBPL-1 (R&D Systems, product no: AF4130, (1:50)), rabbit anti GFP (ThermoFisher, product no: a11122, lot: 1789911 (1:500)), mouse anti NeuN (Millipore, product no: MAB377, lot: 3045564 (1:300)), anti mouse MBP (Millipore, product no: MAB3861, (1:100), rabbit anti GFAP (Dako, product no: Z0334, lot: 00019620 (1:400)), rabbit anti VGAT (Synaptic System, product no: 131002, (1:500)), mouse anti PV (Sigma, product no: P3088, (1:1000)), rabbit anti caspase-3 (Cell Signaling, product no: 9661, (1:100)). For double labelling, some primary antibodies were incubated simultaneously. The next day slides were washed 3 times for 5 minutes in PBS 1X and then incubated for 1 hour at RT in the dark with appropriate fluorophore-conjugated secondary antibodies (Alexa Fluor 488, 546 and 633; 1:1000; Thermo Fisher Scientific. e.g. Cat# A-11006, RRID:AB_2534074; Cat# A-11055, RRID:AB_2534102, Cat# A-11001, RRID:AB_2534069, Cat# A-11008, RRID:AB_143165 and Cat# A-11003, RRID:AB_2534071). In all immunofluorescence staining, nuclei were stained with 4,6 diamine-2-phenylindole (1:25000, DAPI, Roche) in PBS 1X for 1 minute. Sections were mounted with Dako fluorescent mounting medium and subjected to fluorescence and confocal microscope analysis.

Human brain tissue samples were obtained from the Carlo Besta Neurological Institute (Milan, Italy). Research use of human tissue was in accordance with the Declaration of Helsinki (1964– 2008) and the Additional Protocol on the Convention of Human Rights and Biomedicine concerning Biomedical Research (2005). For human tissue, we performed immunohistochemistry for IGFBPL-1. Briefly, brain sections were treated with Xylene for 4 hours, and then rehydrated with decreasing concentrations of ethanol (100, 70, 50 %). Slides were washed in 1X PBS for 5 minutes twice and then incubated in 0.3% H_2_O_2_ in 1X PBS for 10 minutes. To prevent unspecific binding, sections were incubated with blocking solution for 1 hour at RT. The sections were then incubated overnight at +4 °C with the primary antibody rabbit anti IGFBPL-1 pab (Creative Diagnostics, product no: DPABH-0329, (1:100)) diluted in blocking solution in a humidity chamber. The next day the slides were rinsed 3 times for 5 minutes in 1X PBS and then incubated for 1 hour at RT with anti-rabbit biotinylated secondary antibody (Vector Laboratories, product no: BA-1000 (1:500)) diluted in blocking solution. Sections were then rinsed and further incubated with an avidin-biotin complex (ABC reagent, Vector Laboratories product no: PK-6100) for 1 hour at RT. The solution has been prepared 1 hour before use, 10 μl of solution A + 10 μl of solution B in 1 ml 1X PBS. Slides were again washed for 5 minutes 3 times in PBS 1X, incubated in diaminobenzidine (DAB) solution (DAB kit, Vector Laboratories product no: SK-4100) for 1 minute, washed in ddH_2_O and then dehydrated with increasing concentrations of ethanol (50, 70, 100, 100%). Finally, slices were put in xylene for 4 minutes and then mounted with non-aqueous DPX. Staining omitting the primary antibody was always used as negative control.

### Confocal microscopy and image analysis

Confocal (Leica SP5 equipped with 40X objective, Germany) microscopy images were obtained to analyse immunofluorescence staining. Four to five coronal sections were examined for each animal in the SVZ and the other areas of the brain. For neurogenesis quantification, one systematic random series of sections per mice was stained (i.e., for all abovementioned antibodies), from the 14μm thick cryostat coronal sections of the ablated and non-ablated mice, so that sections were spaced at 28 section intervals (280um) from each other. The section series represented a systematic random sample of sections that covered the entire extent of the forebrain.

Cell numbers in the right and left dorsal subventricular zone were quantified manually using Adobe Photoshop CC software on at least 4 sections per mouse containing the SVZ.

For immunohistochemistry staining, images were obtained using Zeiss AxioImager M2m with Nuance FX Multispectral Tissue Imaging System; acquired at 10x and 20x resolution.

### Stereological analysis of Golgi-stained striatal neurons

Mice were anesthetized with ketamine/xylazine (n=3-4 per group), the brains removed and stained using a Rapid GolgiStain™ Kit (FD NeuroTechnologies), and 60μm coronal sections cut. Medium spiny neurons in the striatum close to the SVZ were analysed blinded in the region between Bregma 0 and −1.2 mm. Seven neurons per animal were analysed by using a 40X.

Dendritic branches and spines of the cells were traced using Neurolucida 8.0 (MBF Biosciences, Williston, VT) following the structures through the Z axis. Cell tracings were analysed in Neurolucida Explorer and using the morphometric analysis provided in the Neurolucida® Explorer software package. The morphometric analysis provided in this software package is a 3D Sholl analysis using concentric spheres with a distance of 10 μm between each sphere. The parameters that were analysed included: total dendritic length and number of dendritic intersections. Spine length and spine density was determined using the measuring tool on the StereoInvestigator software (MicroBrightField).

### Electron microscopy

For EM studies ketamine/xylazine anesthetised mice (n= 3/group) were perfused transcardially with 0.9% saline, followed by Karnovsky’s fixative (2% paraformaldehyde and 2.5% glutaraldehyde). The brains were removed and post-fixed in the same fixative overnight. Then, the brains were washed in 0.1 M phosphate buffer. Transverse 200 μm-tick brain sections were cut on a vibratome, post-fixed in 2% osmium for 1 hour, rinsed, dehydrated, and embedded in TAAB resin (TAAB Laboratories, England, UK). Ultrathin sections were mounted onto slot grids for viewing using a LEO 912 transmission electron microscope, as previously described ^59^.

### Discriminative delay-conditioning procedure

Experiments were conducted using 4 identical fully automated Classic Modular Test Conditioning Chambers for Mouse (Med-Associates Product # ENV-307A), for details see Moré et al ^27^ A food-tray was mounted in the centre of the right wall, with an opening located 1 cm above the grid floor and connected to a pellet dispenser through which 14-mg sucrose pellets (Formula P) could be delivered (US). Head entries to the food-tray were detected and recorded by infrared light-beam across the opening. A loudspeaker produced auditory stimuli: two pure sounds at 4-kHz and 9-kHz (CS A and B). Med-PC controlled all the experimental events and recorded the time at which events occurred with 10-ms resolution.

The procedure was repeated for four consecutive days. Two kinds of trials were administered (A+ and B-): both trials started with an Inter Trial Interval (ITI; 160 seconds), followed by a CS (20 seconds duration). The end of the CSs was either paired with food reward (A+) or not (B-). The Med-PC software delivered 30 trials for each trial type (A+ or B-) in a random way for every daily session. Head entries into the food-tray were measured during CSs. Two parameters of mice behaviour were measured: the total number of hits during trials A+ vs. B-, and the timing of the hits during the CSA+ stimulus (hit frequency during the 20 seconds CS presentation, pooled over all trials in a given session).

### Slice preparation and electrophysiology

All procedures were approved by the Italian Ministry of Health and were conducted in accordance with FELASA guidelines and European directives (2010/63/EU). Patch-clamp recordings were performed in sagittal striatal slices. Mice of both sexes (45–60 days of age for young mice, 18 months for aged mice) were anesthetized with an intraperitoneal injection of a mixture of ketamine/xylazine (100 mg/kg and 10 mg/kg, respectively) and perfused transcardially with ice-cold artificial cerebrospinal fluid (ACSF) containing (in mM): 125 NaCl, 3.5 KCl, 1.25 NaH_2_PO_4_, 2 CaCl_2_, 25 NaHCO_3_, 1 MgCl_2_, and 11 D-glucose, saturated with 95% O_2_ 5% CO_2_ (pH 7.3). After decapitation, brains were removed from the skull and 300 μm-thick sagittal slices containing the striatum were cut in ACSF at 4°C using a VT1000S vibratome (Leica Microsystems, Wetzlar, Germany). Slices were then kept in a chamber containing ACSF at 31.5°C for 15 minutes and slowly cooled down and maintained at 26.5°C. Subsequently, individual slices were submerged in a recording chamber mounted on the stage of an upright BX51WI microscope (Olympus, Japan) equipped with differential interference contrast optics (DIC) and an optical filter set for the detection of tdTomato red fluorescent light (Semrock, Rochester, NY, USA). Slices were perfused with ACSF continuously flowing at a rate of 2-3 ml/min at 32°C. Whole-cell patch-clamp recordings were performed in the dorsal striatum using pipettes filled with a solution containing the following (in mM): 30 KH_2_PO_4_, 100 KCl, 2 MgCl_2_, 10 NaCl, 10 HEPES, 0.5 EGTA, 2 Na_2_-ATP, 0.02 Na-GTP, (pH 7.2 adjusted with KOH, tip resistance 6-8Ω). Inter-somatic distances between cells selected for dual recordings were consistently less than 100μm. All recordings were performed using a MultiClamp 700B amplifier interfaced with a PC through a Digidata 1440A (Molecular Devices, Sunnyvale, CA, USA).

#### NPCs application

To study the effect of NPCs application on striatal circuitry, cells (either normal aNPCs, treated with PFA or silenced with shRNA) were collected and resuspended to a concentration of 10^6^ cells/50 μl and gently placed onto the surface of the brain slice at least 30 min before the electrophysiological recordings as already described ^24^.

#### Data acquisition and analysis

Data were acquired using pClamp10 software (Molecular Devices) and analysed with Origin 9.1 (Origin Lab, Northampton, MA, USA). Voltage- and current-clamp traces were sampled at a frequency of 30 kHz and low-pass filtered at 2 kHz. To isolate GABA-receptor dependent IPSCs, the ACSF was added with the AMPA antagonist NBQX (5 μM). Recordings were performed at a holding potential of −80 mV. Spontaneous IPSCs were analysed using pClamp automatic event detector. In dual whole-cell recordings, unitary IPSCs were considered spike-induced when the IPSC onset occurred at a latency of 1-2ms form the peak of the presynaptic spike. Responses were classified as failures when presynaptic spikes induced no IPSCs. To assess synaptic vesicle (SV) readily releasable pools (RRPs) in FSI-MSN pairs, high-frequency stimulus trains (20 Hz, 1.5 s) were applied to the presynaptic FSIs. Cumulative IPSC peak amplitudes were plotted against stimulation time and the linear phase of the dataset was fit with a straight line. Back-extrapolation of the fit line to the Y-axis intercept provided an estimate of the RRP ^60^.

#### Photostimulation of CHR-expressing interneurons

An optogenetic approach was used to selectively stimulate striatal somatostatin expressing (SOM+) interneurons, due to their relative sparseness and long-range connectivity ^19^ which greatly reduces the success rate of dual patch recordings. Optical stimuli were generated by a diode-pumped solid-state laser (wavelength: 473 nm; light power at the source: 100 mW; Shanghai Dream Lasers Technology, Shanghai, China) connected to the epi-illumination port of the microscope through a multi-mode optical fibre. The beam was deflected by a dichroic mirror and conveyed to the slice through a 40x water-immersion objective (spot size: n0.06 mm^2^). The light power measured with an optical power meter at the level of the slice surface was ∼2 mW, yielding a light density value of ∼33 mW/mm^2^. Photostimuli were TTL-triggered using Clampex digital output signals.

#### Drugs

The following drugs were obtained from HelloBio (Bristol, UK): NBQX disodium salt (HB0443) and SR95531 hydrobromide (*Gabazine*, HB0901). IGF-1 (PeproTech; 100-11) was added to the chamber at 26.5°C to a final concentration of 10nM at least 30 minutes before the electrophysiological recordings.

### Synapses segmentation and analysis

Synapses were segmented by a custom routine as previously reported ^61^. A collection of regions of interest (ROI) was run on vGAT fluorescence channel, synapses segmentation was based on fluorescence, shape, size and Voronoid tassellation. A fluorescence threshold for puncta inclusion/exclusion was set as follows: mean background fluorescence was calculated, for each field analysed, from ROIs devoid of fluorescent puncta. All the segmented objects with mean fluorescence > mean background +2SD were considered as synapses, the remaining were excluded. Synapses were considered positive for tdTomato when average fluorescence was higher than the background tdTomato fluorescence calculated as described above. Double positive synapses were than normalized over total perisomatic boutons (vGAT positive only) and represented as percentages.

### RNA extraction

Mice were sacrificed under deep anaesthesia with tribromoethanol (Avertin) and transcardially perfused with cold saline solution with EDTA. Brains were rapidly removed and SVZ were dissected on ice and stored at −80°C until RNA extraction.

Total RNA was isolated using the RNeasy MiniKit (#74104; Qiagen) according to the manufacturer’s instructions, including DNAse digestion. At the end, RNA samples were eluted from columns using 30 μl of RNase-free water and their concentrations were determined or spectrophotometrically by A260 (Nanodrop–ND1000) or using a 2100 Bioanalyzer (Agilent) resulting in RNA integrity number ≥8.

### Gene expression analysis

Semi quantitative RT-PCR was performed using pre-developed TaqmanTM Assay Reagents on a Quantstudio 3 Real-Time PCR System (Thermofisher) according to manufacturers’ protocol.

The cDNA was synthesized from 500 ng of total RNA using the ThermoScript IV RT-PCR System (Thermofisher) according to the manufacturer instructions. Two-hundred ng of cDNA were used for RT-PCR using pre-designed Taqman® Gene Expression Assays (Thermofisher). RT-PCRs were performed using the following specific assays (all Thermofisher): *Igfbpl-1* (Mm01342060_m1), *Igf-1r* (Mm00802831_m1), *Dcx* (Mm00438400_m1), *Dlx2* (Mm00438427_m1), *Igfbpl-1* (Hs01390103_m1).

Data were collected with instrument spectral compensations by the Thermofisher Connect™ software and analysed using the threshold cycle (CT) relative quantification method. In this study, GAPDH was used exclusively as housekeeping gene.

### RNA-sequencing

For SVZ tissue libraries were prepared using SMART-Seq® v4 Ultra® Low Input RNA Kit for Sequencing (Takara Bio USA) performing 10 cycles of amplification, according to the manufacturer’s instructions, while for the striatum tissue libraries were prepared using QuantSeq 3’ mRNA-Seq Library Prep Kit FWD for Illumina (Laxogene).

Sequencing was performed on a NextSeq 500 machine (Illumina, San Diego, CA) obtaining 30 million single end reads per sample on average.

Reads were trimmed using Trimmomatic, version 0.32, in order to remove adapters and to exclude low-quality reads from the analysis. The remaining reads were then aligned to the reference genome mm10, Gencode version M16, using STAR aligner, version 2.5.3a. The feature Counts function from Rsubread package (v 1.16) was used to assign reads to the corresponding genes. Only genes with a CPM (Counts per million) value higher than 1 in at least three samples were retained. Gene expression read counts were exported and analysed in R environment (v. 3.1.1) to identify differentially expressed genes (DEGs), using the limma Bioconductor library ^62^. DEGs were identified as those genes that changed at least 2-fold and had a nominal pvalue lower than 0.01. Pre-ranked Gene Set Enrichment Analysis (GSEA) ^63^ was performed considering all the expressed genes. The gene-sets included in the GSEA analyses were obtained from Canonical Pathways, Hallmark and the Gene Ontology (GO) collections as they are reported in the MSigDB database (https://www.gsea-msigdb.org/gsea/msigdb/index.jsp).

### Assay for transposase-accessible chromatin using sequencing (ATAC-seq)

Single-cell ATAC-seq was performed on Chromium platform (10X Genomics) using “Chromium Single Cell ATAC Reagent Kit” V1 Chemistry (manual version CG000168 Rev C), and “Nuclei Isolation for Single Cell ATAC Sequencing” (manual version CG000169 Rev B) protocols. In-house produced Tn5 protein with modified Tn5ME-A sequence (manuscript under revision) was used instead of the “ATAC Enzyme” (10X Genomics). Nuclei suspension was prepared in order to recover and analyze 5,000 nuclei. Libraries were sequenced on Novaseq6000 platform (Illumina) with 2×50 bp read length; a custom Read 1 primer was added to the standard Illumina mixture (5’-TCGTCGGCAGCGTCTCCGATCT-3’) (manuscript under revision).

Reads were demultiplexed using cell ranger-atac (v1.0.1). Identification of cell barcodes was performed using umitools (v1.0.1) ^64^ using R2 as input. Read tags were aligned to hg38 reference genome using bwa mem v0.7.12 [arXiv:1303.3997 (q-bio.GN)]. Gene activities were calculated as per cell coverage over the gene body interval extended to 2kb upstream the TSS, using gencode v30 as gene model ^65^. Data were processed using scanpy v1.4.6 ^66^.

Pseudobulk tracks were generated using pyGenomeTracks ^67^.

### In situ-hybridization

In situ hybridization was performed as previously described ^68,69^. Briefly, 14 μm-thick brain sections were post-fixed 15 min in 4% paraformaldehyde, then washed three times in PBS. Slides were incubated in 0.5 mg/ml of Proteinase K (Roche) in 100 mM Tris-HCl (pH 8), 50 mM EDTA for 10 min at 30°C. This was followed by 15 min in 4% Paraformaldehyde. Slices were then washed three times in PBS then washed in H_2_O. Sections were incubated in triethanolamine (Merk, Germany) 0.1 M (pH 8) for 5 min, then 400 ml of acetic anhydride (Sigma) was added two times for 5 min each. Finally, sections were rinsed in H_2_O for 2 min and air-dried. Hybridization was performed overnight at 60°C with a-UTP riboprobes at a concentration of 100 ng/ul. The following day, sections were rinsed in SSC 5 X for 5 min then washed in formamide 50% (Sigma)-SSC 2 X for 30 min at 60°C. Then slides were incubated in ribonuclease-A (Roche) 20 mg/ml in 0.5 M NaCl, 10 mM Tris-HCl (pH 8), 5 mM EDTA 20 min at 37°C. Sections were washed in formamide 50% SSC 2X for 30 min at 60°C then slides were rinsed two times in SSC 2X. After that, the sections were blocked in B1 buffer [150 mM NaCl, 100 mM Tris-HCl (pH 8)] and 10% of foetal bovine serum for 1 hour. Finally, the sections were incubated overnight at +4 °C with the antibody anti-digoxigenin-AP Fab fragment (#11093274910, Roche, 1:1000) diluted in blocking solution in a humidity chamber. The next day, sections were washed for 10 minutes 2 times with buffer B3 [100 mM NaCl, 100 mM Tris-HCl (pH 9.5), MgCl_2_ 50 mM] and incubated in 5-bromo-4-chloro-3-indolyl-phosphate (BCIP) (Roche) and 4-nitro blue tetrazolium chloride (NBT) (Roche) overnight. The following probes were used: mouse *Igf-1r* riboprobe (gift from Giacomo Consalez, San Raffaele Hospital, Milan, Italy). Microphotographs of sections were digitalized in dark field light microscopy (Olympus BX51, and 46 objective) by using a CCD camera (Leica). To confirm the specificity of the different RNA probes, sense strand RNA probes (showing no signal) were used as negative controls. As positive control for the technic we use *plp* riboprobe (present in the laboratory.

### Mouse and human neural stem cell cultures: generation and maintenance

#### Mouse NPCs

Adult C57B/6 mice (6 to 8 weeks old, 18-20 g Charles River) were anaesthetized by an intraperitoneal injection of tribromoethanol and killed by decapitation. The parietal bones were cut in a cranial-to-caudal way using microsurgery scissors and the brain was removed and positioned in a Petri dish (Corning Costar) containing sterile PBS. Brain coronal sections were taken 2 mm from the anterior pole of the brain, excluding the optic tracts and 3 mm posterior to the previous cut. The SVZ of the lateral ventricles was isolated from the coronal section using iridectomy scissors. Tissues derived from at least two mice were pooled to generate each culture. Dissected tissues were transferred to a 15 ml tube with digestion medium [EBSS 1X (Gibco), Cysteine (Sigma) 200 mg/L, EDTA (Sigma) 200mg/L, Papain (Worthington) 2U/ml], and incubated for 45 min at 37°C on a rocking platform, as previously described ^59^. At the end of the incubation, the tube was centrifuged at 200 g for 12 minutes, the supernatant was removed, and the pellet was mechanically disaggregated with 2 ml of EBSS. The pellet was centrifuged again at 200 g for 12 minutes and then dissociated with a 200μl pipette and plated in Neurocult proliferation medium (Stem cell Technology, BC, CA) supplemented with EGF (20 ng/ml) and FGF2 (10 ng/ml). Cells were plated in a 25 cm^2^ flask (Corning Costar) and incubated at 37°C in an atmosphere of 5% CO_2_. After approximately one week, a small percentage of the isolated cells begun to proliferate, giving rise to small cellular aggregates, which are similar to rounded spheres (called neurospheres) and which grow in suspension. When the neurospheres reached the necessary dimension (about ∅ ≥ 150-200 μm diameter), the cells were harvested in a 15 ml tube (Falcon) using a sterile pipette and centrifuged at 200 g for 12 minutes. The supernatant was then removed, 200 μl of Accumax (Sigma) was added and the tube was incubated at 37°C for 10 minutes. The cells were dissociated, counted and plated at the density of 8000 cells/cm^2^. This procedure was repeated at each passage of dissociation.

#### Human foetal NPCs

Permission to use human foetal CNS tissue was granted by the ethical committee of the San Raffaele Hospital (approval on 09/06/2016). Tissue procurement was in accordance with the declaration of Helsinki and in agreement with the ethical guidelines of the European Network for Transplantation (NECTAR). The BI-0194-008 is a non-immortalized human NPC line obtained from the diencephalic and telencephalic regions of a single human caucasian male foetus at 10–12 weeks gestational age after pregnancy termination. The fetus was provided by Banca Italiana del Cordone Ombelicale Fondazione IRCCS Ca’ Granda Ospedale Maggiore Policlinico, Milan, Italy.

Briefly, fresh human foetal brain tissue was chopped mechanically and dissociated with Trypsin (LONZA; BE17-161E) 1:5 in growth medium for 5-10 min at 37°C, 5% O2 e 5% CO2. Then, it was washed with 10% Australian FBS in fresh medium and centrifuged 15 min at 200 g. Finally, the cells were plated in T25 flask in NeuroCult-XF Proliferation Medium (STEMCELL Technologies; cat.# 05760) with rh-EGF (Provitro; cat.# 1325 9510 00) and rh-bFGF (Provitro; cat.# 1370 9505 00) at the final concentration of 20 ng/ml, at 37°C, 5% O2 e 5% CO2.

After 15-25 days, enzymatic (Accumax®; Sigma) dissociation of neurospheres was performed at 37°C for 3-5 minutes, of neurospheres was performed and the cells were re-plated at clonal density (20-25000 cells/cm2) in NeuroCult-XF Proliferation Medium completed as mentioned above.

#### Human induced-pluripotent stem cells (iPS) derived NPCs

Permission to generate induced-pluripotent stem cells (iPS) from MS patients’ fibroblasts was granted by the ethical committee of the San Raffaele Hospital (BIOBANCA-INSPE, approved on March 9^th^, 2017). To avoid genetic biases, fibroblasts were isolated from skin biopsies from a pair of monozygotic twins discordant for MS (females, age 34 at time of tissue donation, 3 years from disease onset in the affected twin). Fibroblasts were reprogrammed into iPSCs by using the replication-incompetent Sendai virus kit (Invitrogen) according to manufacturer’s instructions. In brief, colonies of iPSCs were detached from mouse embryonic fibroblasts by treatment with 1 mg/mL collagenase IV. After sedimentation, cells were resuspended in human embryonic stem cell (hESC) medium without bFGF2 supplemented with 1 µM dorsomorphin (Tocris), 3 µM CHIR99021 (Axon Medchem), 10 µM SB-431542 (Ascent Scientific) and 0.5 µM purmorphamine (Alexis). Embryoid bodies (EBs) were formed by culturing cells in non-tissue-culture petri dishes (Corning). On day 2 medium was changed to N2B27 medium containing equal parts of neurobasal (Invitrogen) and DMEM-F12 medium (Invitrogen) with 1:100 B27 supplement lacking vitamin A (Invitrogen), 1:200 N2 supplement (Invitrogen), 1% penicillin/streptomycin/glutamine (PSG) and the same small molecules mentioned before. After two additional days, dorsomorphin and SB-431542 were withdrawn, while ascorbic acid (AA; 150 µM) was added to the medium. On day 6, EBs were cut into smaller pieces and plated onto matrigel (Matrigel Growth-factor-reduced, Corning) coated 12-well plates. For passaging, NPCs were treated with accutase. After 3 passages purmorphamine was replaced by 0.5 µM SAG (Cayman Chemical) (NPC medium).

### Immunofluorescence on mouse and human cells

For *in vitro* analysis, mouse neurosphere suspensions and 250.000 of iPS-derived human NPCs were plated on MATRIGEL-coated (Becton Dickinson Labware) round 12-mm coverslips in a 24-well plate in 1 ml of Neurocult medium plus proliferation supplement.

Neurospheres were fixed with 4% PFA after 2 hours, while iPS-derived human NPCs after 3 days, at room temperature (RT) for 10 minutes, then rinsed three times with PBS 1X, and then incubated for 60 min at RT with a blocking solution [PBS 1x + 10% normal goat serum (NGS)]. For intracellular staining, the same blocking solution as above, plus 0.1% Triton X-100, was used. Then fixed cells were incubated for 2 further hours at RT with an appropriate primary antibody diluted in PBS 1X + 1% NGS. Primary antibodies used: purified goat anti mouse IGFBPL-1 (R&D Systems, product no: AF4130, (1:50)), rabbit anti mouse Olig2 (Millipore, product no: MABN50, lot: 211F1.1 (1:200)), mouse anti mouse Nestin (Millipore, product no: MAB353, (1:100)), rabbit anti mouse Sox2 (Abcam, product no: ab69893, (1:200)), rabbit anti human IGFBPL-1 pab (Creative Diagnostics, product no: DPABH-0329, (1:100)), and mouse anti human Nestin (Millipore, product no: MAB5326, (1:200)). Cells were then washed three times in 1X PBS and then incubated for 45 minutes with the appropriate fluorescent secondary antibodies (1:1000, AlexaFluo). The nuclei were stained with 4,6 diamine-2-fenilindole (1 μg/ml, DAPI, Roche). Cells were then washed and mounted with Fluorescent mounting medium (Dako).

Light (Olympus, BX51, Japan) and confocal (Leica, SP5 equipped with 40X and 63X objectives, Germany) microscopy images were obtained to analyse cell stainings. Image analyses was performed using Adobe Photoshop CS4 software (Adobe Systems Incorporated) or ImageJ (NIH software).

### Elisa assay

Mice were killed by cervical dislocation, and the brains were rapidly removed after decapitation and the SVZ and striatum were dissected. The samples were homogenized in 200 μl of homogenization buffer (10 mM Tris HCl pH7.4, 260 mM sucrose and cocktail of protease inhibitors). The homogenate was centrifugated at 1000 X g at 4°C for 10 min to remove nuclei and cell debris. Total proteins were quantified by using BCA kit (Pierce, Thermo Scientific IL 61101 USA). ELISA for IGF-1 (DuoSet ELISA kit, R&D Systems, product no: DY791) was performed according to the manufacturer’s instructions.

### In-vivo local field potentials (LFPs) recordings and analysis

Teflon-insulated stainless-steel wires (Ø 150 μm) were implanted in the right and left striatum and primary sensory cortices (S1). Electrodes were built with a single wire connected to a pin. Under deep sevofluorane anaesthesia, electrodes were stereotaxically implanted in the striatum and S1, according to the following coordinates, in mm: striatum, AP = 0.5, L= 1.5, V=-2.9; S1, AP= −1.0, L=3.0, V=-1.8. A silver wire over the cerebellum was used as reference and ground. All implants were secured using Ketacem cement. After surgery, mice were allowed to recover for 6-7 days before testing. All recordings were performed inside a customized Faraday chamber and lasted 10 minutes each, within the home cage. LFPs were recorded and initially digitized at 1 kHz then stored on a hard drive for offline analysis. LFPs epochs were visually examined and power spectra of artefact-free segments were computed using fast Fourier transforms by using the commercial software NeuroExplorer (Plexon). Mean power spectra were divided into frequency bands: 2-7 Hz, 7.01-12 Hz, 12.01-20 Hz, 20.01-30 Hz, 30.01-70 Hz and 70.01-12o Hz ^70^. Three epochs of 10 sec CS were analysed for each animal and averaged.

Interhemispheric coherence between LFP channels was measured by magnitude squared coherence (MSC), using the function ms cohere in Matlab signal toolbox, which is a coherence estimate of the input signals x and y by using Welch’s averaged, modified periodogram method. The MSC estimate is a function of frequency with values between 0 and 1 and indicates how well x corresponds to y at each frequency. The MSC estimate is calculated over the frequency range of 0.5 – 120 Hz for each animal. Statistics were performed on the average difference in coherence within the frequency bands of interest ^71^.

### MRI human experiment

We retrospectively evaluated MRI scans acquired from 97 patients with MS and 43 age- and sex-matched healthy controls. For all participants, exclusion criteria were age > 50 years, history of drug or alcohol abuse, neurologic disorders (excluding MS in patients), psychiatric comorbidities, history of head trauma and contraindications to MRI. The age range was chosen to minimize possible confounding effects of unknown chronic small vessel disorders of the brain. Patients had a diagnosis of clinically isolated syndrome (n=4) or MS (n=93), according to 2017 revision of McDonald criteria.^72^ Ethical approval was received from the local ethical standards committee of the San Raffaele Scientific Institute, Milan (Italy), and written informed consent was obtained from all participants at the time of data acquisition.

On the same day of MRI, the Symbol Digit Modalities Test (SDMT) was administered to patients and results were education-corrected according to normative data to obtain Z-scores.^40^

Using a 3.0 Tesla Philips Ingenia CX scanner with a dS Head 32-channel receiver coil and standardized procedures for subjects positioning, the following brain MRI protocol was acquired: 1) variable flip angle 3D T2-weighted fluid-attenuated inversion recovery (FLAIR) turbo spin echo (repetition time [TR]= 4800 ms; echo time [TE]= 270 ms; inversion time [TI]= 1650 ms; matrix size = 256 × 256; field of view [FOV]= 256 × 256 mm^2^; echo train length [ETL]= 167; 192 contiguous 1 mm-thick sagittal slices); 2) 3D T1-weighted turbo field echo (TR= 7 ms; TE= 3.2 ms; TI= 1000 ms; flip angle= 8°; matrix size= 256 × 256; FOV= 256 × 256 mm^2^; 204 contiguous 1 mm-thick sagittal slices) and 3) diffusion-weighted pulsed-gradient spin-echo single-shot echo-planar (TR= 5900 ms, TE= 78 ms; matrix size= 112 × 85; FOV= 240 × 233 mm^2^; 56 contiguous 2.3 mm-thick slices; number of excitations [NEX]= 1) with diffusion-weighting (b-factor= 700/1000/2850 s/mm^2^) applied along 6/30/60 noncollinear directions and ten b=0 volumes distributed along the acquisition, plus three additional b=0 volumes with reversed polarity of gradients for distortion correction were acquired.

T2-hyperintense brain lesions masks were automatically segmented using the FLAIR and 3D T1-weighted scans as input images^73^ and corresponding volumes were calculated.

After 3D T1-weighted images lesion refilling, head-size-normalized volumes of the entire brain, grey matter, white matter and deep grey matter nuclei (caudate, pallidum, putamen, thalamus and hippocampus) were segmented and measured using FSL SIENAx software and FMRIB’s Integrated Registration and Segmentation Tool (FIRST) pipeline (FMRIB, Oxford, UK).^74,75^ On the same refilled 3D T1-weighted images, average and lobar (frontal, temporal, parietal, occipital and cingulate) cortical thickness was obtained running the FreeSurfer 6.0 software suite (http://surfer.nmr.mgh.harvard.edu/).^76^

Pre-processing of diffusion-weighted images included correction for off-resonance and eddy current induced distortions, as well as for movements using the Eddy tool within the FSL library.^77^

Then, the diffusion tensor was estimated from the two lower shells by linear regression^78^ using FSL software (FMRIB, Oxford, UK).

Fractional anisotropy and mean diffusivity maps were derived and measured within lesions and in the normal appearing tissue, defined as those voxels not involved by T2-hyperintense lesions.

A SVZ mask was segmented on 3D T1-weighted images in the standard space (Montreal Neurological Institute space) according to anatomical references based on lateral ventricles and caudate segmentation. ^39^

SVZ and T2-hyperintense lesion masks were registered on native 3D T1-weighted and diffusion weighted images to obtain the percentage of lesioned SVZ (SVZ percentage lesion volume) and intralesional/normal appearing tissue measures of microstructural integrity.

Age-, sex- and phenotypes-adjusted linear models, partial correlations and stepwise multiple linear regressions were used to assess SVZ damage and to identify predictors of caudate and SDMT z-scores.

For partial correlations and stepwise multiple linear regressions, all the following measures of SVZ and brain damage were included: SVZ percentage lesion volume, fractional anisotropy in the normal appearing tissue of SVZ, intralesional fractional anisotropy in the SVZ, mean diffusivity in the normal appearing tissue of SVZ, intralesional mean diffusivity in the SVZ, brain volume, grey matter volume, white matter volume, logarithm of T2-hyperintense lesion volume, deep grey matter volume (excluding the caudate), caudate volume, white matter fractional anisotropy, white matter mean diffusivity, mean cortical thickness and cortical thickness of the frontal, parietal, temporal, occipital and cingulate lobes.

### Statistical analysis

For statistical analyses, we used a standard software package (GraphPad Prism version 7.00). Data were evaluated by unpaired two-tailed t-test (for comparisons between 2 groups) or by one-way ANOVA followed by post hoc analysis (for comparison among 3 groups) as indicated in the figure legends. The significance level was established at p=0.05. A post hoc analysis was performed using Bonferroni correction. For local field potential experiments Kruskal-Wallis was used followed by Dunn’s correction. For patch clamp experiments we used Unpaired two-tailed t-test, One-way ANOVA followed by Tukey post-test and Z-Test for 2 Population Proportions. For behavioural experiments, one-way ANOVA was used to compare the frequency of nose pokes during the presentation of the CS among the different genotypes, whereas the time courses of the poking frequencies were compared by transforming them into cumulative probability distributions (pokes vs. time) and contrasting the distributions obtained in different genotypes using the Kolmogorov-Smirnov test.

## Acknowledgements

We thank the Advanced Light and Electron Microscopy BioImaging Center of the San Raffaele Scientific Institute; Dong Feng Chen’s laboratory for kindly providing *Igfbpl-1*^-/-^ mice; Giorgio Giaccone and Fabrizio Tagliavini for kindly providing human SVZ tissue. This work was supported by Progressive MS Alliance (collaborative research network PA-1604-08492 (BRAVEinMS) to GM and TK), the Italian Multiple Sclerosis Foundation ((FISM) Project No. ‘Neural Stem Cells in MS’ to GM), and FRRB (Project PLANGECELL to GM).

