## Supplementary Figures for "Neural precursor cells contribute to decision-making by tuning striatal connectivity via secretion of IGFBPL-1"

**Supplementary Fig. 1 Neurogenesis and striatal MSN morphology are rescued in SVZ-eNPC ablated mice**

**a,** Representative images and quantification of eNPC in the SVZ one month after the end of GCV treatment in NestinTK$^-$GCV$^+$ and NestinTK$^+$GCV$^+$ mice: neuroblasts were labelled for doublecortin (DCX, in green) and transient amplifying cells for BrdU (in red). Nuclei in blue were counterstained by DAPI. The lateral ventricle is denoted as LV. Scale bar: 50 µm. n= 3 mice per group. **b, c,** Sholl analysis comparing tracings of medium spiny neurons from NestinTK$^-$GCV$^+$ wash out and NestinTK$^+$GCV$^+$ wash out n=6-8 neurons per mice for a total of 3 mice per group. Data are presented as number of dendritic intersections **(b)**, and total dendritic length plotted at increasing distance from the cell body **(c)**. **d, e,** Striatal MSNs spine length **(d)**, and spine density **(e)** in NestinTK$^-$GCV$^+$ wash out compared to NestinTK$^+$GCV$^+$ wash out mice. Values represent the mean ± SEM.

**Supplementary Fig. 2 The SVZ ablation causes persistent increase in low-frequencies coherence between the right and left striatum**

**a,** Schematic representation of the implanted electrodes (striata in red and somatosensory cortices in blue) and interhemispheric coherence analysis of LFP signal, showing higher striatum-striatum synchrony but not S1-S1**. b, c.** The interhemispheric coherence in the NestinTK$^+$ mice before the treatment with GCV (NestinTK$^+$GCV$^-$), after 4 weeks of GCV treatment (NestinTK$^+$GCV$^+$) and one month after the end of GCV administration (NestinTK$^+$GCV$^+$ wash out) from 2 to 200 Hz between right and left striatum (n = 14 for NestinTK$^+$GCV$^-$, n=13 for NestinTK$^+$GCV$^+$ and n=9 NestinTK$^+$GCV$^+$ in the recovery phase) in a and between the right and left somato-sensorial cortex (n = 6 for NestinTK$^+$GCV$^-$, n=6 for NestinTK$^+$GCV$^+$ and n=9 NestinTK$^+$GCV$^+$ in the recovery phase) in **b**. Values represent means ± SEM. In **a**, for 2-7 Hz ## p= 0.0086, for 7-12 Hz *p=0.0281, ###p=0.0039, for 12-20 Hz *p=0.0358, #p=0.0144; in **b**, for 20-30 Hz ##p=0.0084, for 30-70 Hz *p=0.0315, #p=0.0341. Unpaired Kruskal-Wallis test followed by Dunn's correction was used.

**Supplementary Fig. 3 NPC-ablation does not affect cell-to-cell inhibitory GABAergic transmission in striatal MSN pairs**

**a,** Examples of unitary IPSCs evoked in an MSN in response to individual APs elicited in a synaptically connected MSN. Both successful IPSCs and failures are shown. The drawing represents the recording configuration. **b,** Connectivity rates for MSN-MSN pairs in NestinTK$^+$GCV$^-$ and NestinTK$^+$GCV$^+$ slices. **c,** Summary of average amplitudes of unitary IPSCs in MSN-MSN pairs from NestinTK$^+$GCV$^-$ (n =11 cells in 5 mice) and NestinTK$^+$GCV$^+$ (n =11 cells in 5 mice) slices. Values represent medians, Mann Whitney U test. **d**, Summary histogram comparing reliability of MSN-MSN GABAergic synapses (measured as % of MSNs displaying at least one failure).

**Supplementary Fig. 4 GABAergic transmission mediated by somatostatin (SOM)-expressing interneurons was not impaired after NPC ablation**

**a,** Left, schematic cartoon showing the experimental approach used to optogenetically stimulate striatal SOM interneurons. Right, example of a unitary IPSC and a failure evoked in a MSN in response to optical activation (represented by the blue trace on bottom) of striatal SOM interneurons. **b,** Summaries of connectivity rates of MSNs. n=3 mice for SomChR-NestinTK$^-$GCV$^+$ and n=5 mice for SomChR-NestinTK$^+$GCV$^+$. **c,** Summary plots with unitary parameters (peak amplitude and failure rates) for SOM interneuron-mediated IPSCs in SomChR-NestinTK$^-$GCV$^+$ (n=3) and SomChR-NestinTK$^+$GCV$^+$ mice (n=5). **d,** Summaries of percentages of MSNs showing failures in response to optogenetic stimulation of SOM interneurons 15 out of 22 (68%) and 19 out of 27 (70%) MSNs were found connected with at least one SOM interneuron in SomChR-NestinTK$^-$GCV$^+$ (n=3 mice) and SomChR-NestinTK$^+$GCV$^+$ (n=5 mice) slices respectively. Of these, 46% (7/15) and 53% (10/19) displayed at least one failure.

**Supplementary Fig. 5 SVZ-eNPC ablation causes presynaptic defects in striatal GABAergic terminals**

**a,** Examples of IPSC trains evoked in MSNs in response to extracellular, high-frequency local stimulation (50 Hz, 1s). **b,** Cumulative plots of IPSC peak amplitudes recorded in MSNs from PVCretDt-NestinTK$^-$GCV$^+$ and PVCretDt-NestinTK$^+$GCV$^+$ mice. The linear portion of the cumulative IPSC amplitude distribution was fit with a straight line that was prolonged to intercept the Y-axis, in order to obtain an estimated value of the vesicular readily releasable pool (RRP). **c,** Summary plot of RRP sizes in MSNs from PVCretDt-NestinTK$^-$GCV$^+$ (n=11 pairs in 8 mice) and PVCretDt-NestinTK$^+$GCV$^+$ (n=15 pairs in 12 mice). Values represent means ± SEM. unpaired two-tailed t-test *p=0.048. **d,** Representative confocal microscopy images showing coronal brain sections of the striatum in PVCretDt-NestinTK$^-$GCV$^+$ and PVCretDt-NestinTK$^+$GCV$^+$ mice. Scale bar: 10 μm. **e,** Quantification of presynaptic terminals expressing both PV and vGAT, normalized over the total of vGAT positive presynpases in PVCretDt-NestinTK$^-$GCV$^+$ and PVCretDt-NestinTK$^+$GCV$^+$ mice. Values represent means ± SEM. *p=0.0362; Mann-Whitney U test. n=9 for PVCretDt-NestinTK$^-$GCV$^{+,}$ n=12 for PVCretDt-NestinTK$^+$GCV$^+$ mice. Scale bar: 5 μm.

**Supplementary Fig. 6 Aged mice have reduced neurogenesis and neurophysiological striatal defects**

**a,** Quantification of neurogenesis in young and aged mice. Neuroblasts were labelled for doublecortin (DCX) and transient amplifying cells for BrdU. Nuclei in blue were counterstained by DAPI. The lateral ventricle is denoted as LV. Values represent means ± SEM. **p=0.001, ****p=$2.5*10^{-5}$ Unpaired two tailed t-test, n= 4 young mice and n=6 aged mice. Scale bar: 50 µm. **b,** PCR quantification for *Dcx* and *Dlx2* in young and aged mice. Values represent means ± SEM. ****p=$2.6*10^{-5}$, ***p=0.0005 unpaired two tailed t-test, n= 4 young mice and n=6 aged mice. **c,** Left spontaneous IPSCs recorded in a control young group (n=5 mice) and aged (n=4) C57BL/6 mice. Right summary plot with average sIPSC frequency and amplitude. Values represent means ± SEM. ***p=0.0008, ****p<0.0001, unpaired two tailed t-test. **d,** Histograms summarizing synaptic reliability of FSI-MSN GABAergic connectivity in our reference young vs. aged mice. Z-Test for 2 Population Proportions; *p=0.0046. The number of MSNs showing at least one failure was: 2 out of 15 (13%) in young mice (n=8) and 9 out of 14 (64%) in old mice (n=4). **e,** Quantitative PCR of *Igfbpl-1* and representative confocal images of the SVZ of young and aged mice labelled for IGFBPL-1 in green and NeuN in red. Nuclei in blue were stained by DAPI. The lateral ventricle is denoted as LV. Values represent means ± SEM. ***p=0.0003, unpaired two tailed t-test, n= 3 young mice and n=6 aged mice. Scale bar: 50 µm.

**Supplementary Fig. 7 Gene expression analysis in the striatum of SVZ-eNPC ablated mice**

**a,** Workflow of RNA-seq gene expression analysis in control and NestinTK$^{+}$GCV$^{+}$ treated mice. **b,** Heatmap of hierchical clustering of expression values of 22 down regulated genes and 162 up regulated genes with ≥2-fold change in control and NestinTK$^{+}$GCV$^{+}$ mice. (blue means downregulation, red upregulation) n=5 mice per group. **c,** Violin plot of *Igf-1* pathway average expression in control mice compared to NestinTK$^{+}$GCV$^{+}$ mice. *Igf-1* pathway was retrieved from Reactom Database. (Pre-ranked GSEA: FDR= 0.03, NES= -1.82). **d,** Representative images of the *in situ* hybridization for *Igf-1r* at the level of the striatum in control mice. The inset shows a cortex staining as positive control (PC). **e**, The graph shows the quantification obtained by rt-PCR for *Igf-1r* in the striatum of the control and NestinTK$^{+}$GCV$^{+}$ mice. Values represent the mean ± SEM. **p= 0.0075, unpaired two-tailed t-test. n=12 ctrl mice and n=6 NestinTK$^{+}$GCV$^{+}$ mice. Scale bar: 100 µm. **f,** Quantification of IGF1 protein by Elisa in the SVZ of NestinTK$^{+}$GCV$^{+}$ and NestinTK $^{+}$GCV$^{+}$ mice (n=3 mice per group). Values represent means ± SEM ****p=$1.9*10^{-7}$ unpaired two-tailed t-test.

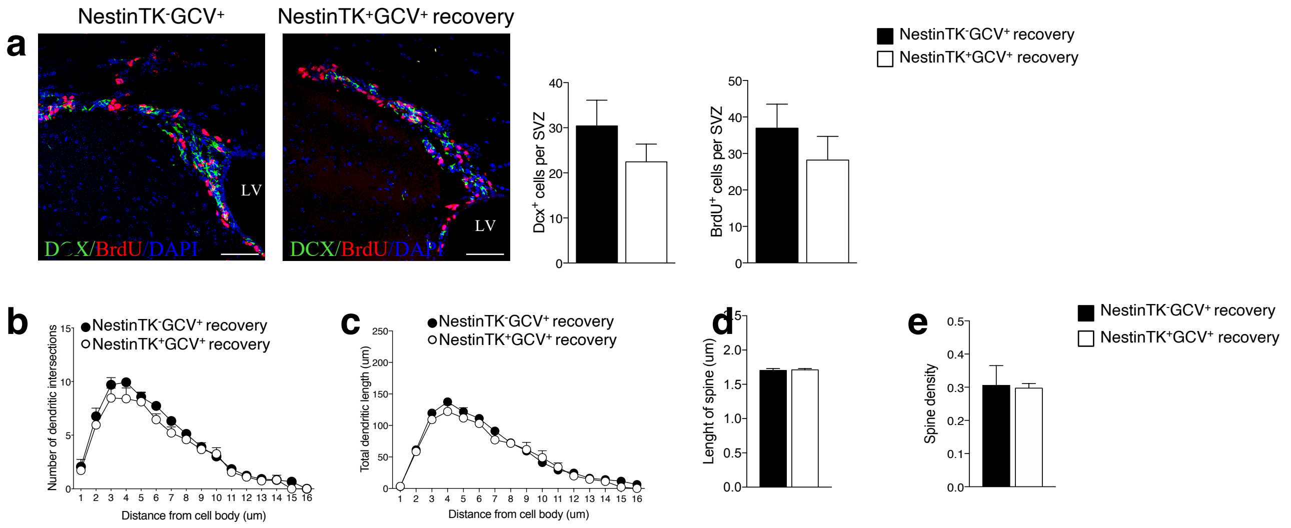
a
NestinTK-GCV+
NestinTK+GCV+ recovery
LV
DCX/BrdU/DAPI
Dcx+ cells per SVZ
BrdU+ cells per SVZ
NestinTK-GCV+ recovery
NestinTK+GCV+ recovery
b
Number of dendritic intersections
Distance from cell body (um)
c
Total dendritic length (um)
d
Lenght of spine (um)
e
Spine density

Figure S1

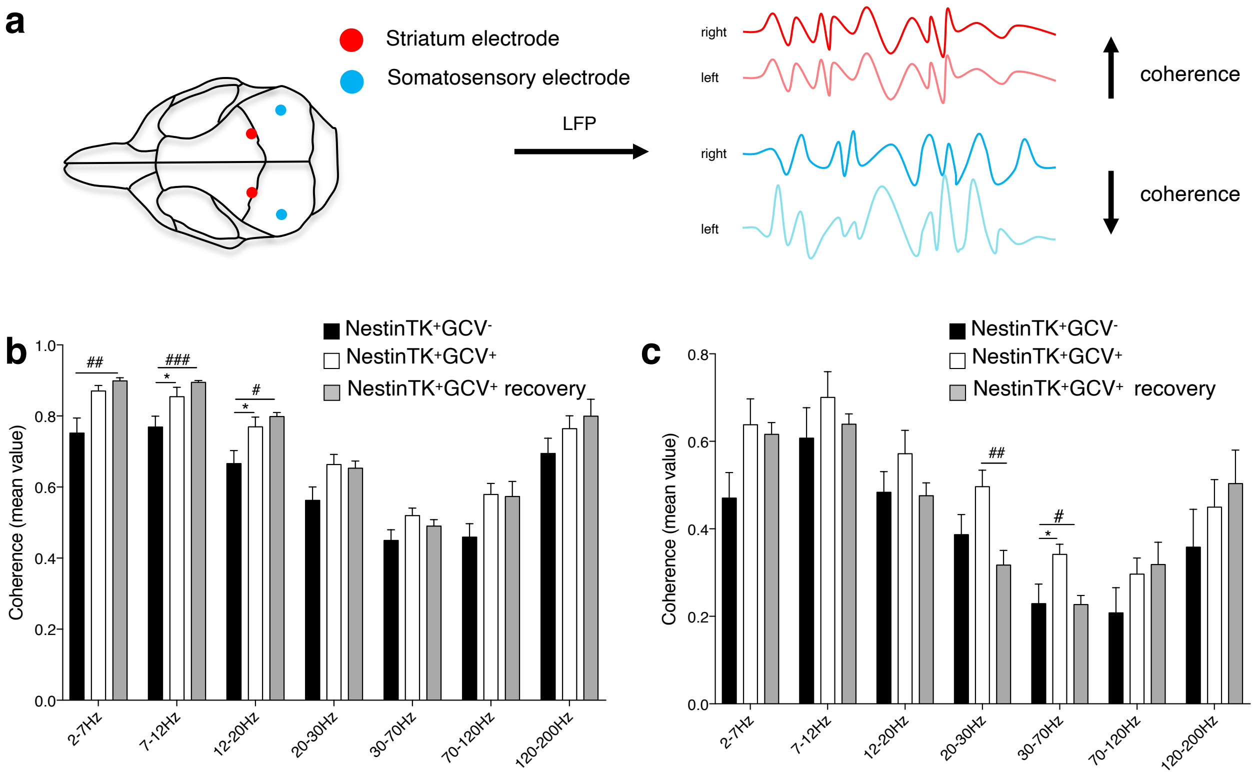
a
Striatum electrode
Somatosensory electrode
LFP
right
left
right
left
coherence
coherence
b
NestinTK+GCV-
NestinTK+GCV+
NestinTK+GCV+ recovery
Coherence (mean value)
1.0
0.8
0.6
0.4
0.2
0.0
##
###
*
#
*
2-7Hz
7-12Hz
12-20Hz
20-30Hz
30-70Hz
70-120Hz
120-200Hz
c
NestinTK+GCV-
NestinTK+GCV+
NestinTK+GCV+ recovery
Coherence (mean value)
0.8
0.6
0.4
0.2
0.0
##
#
*
2-7Hz
7-12Hz
12-20Hz
20-30Hz
30-70Hz
70-120Hz
120-200Hz

Figure S2

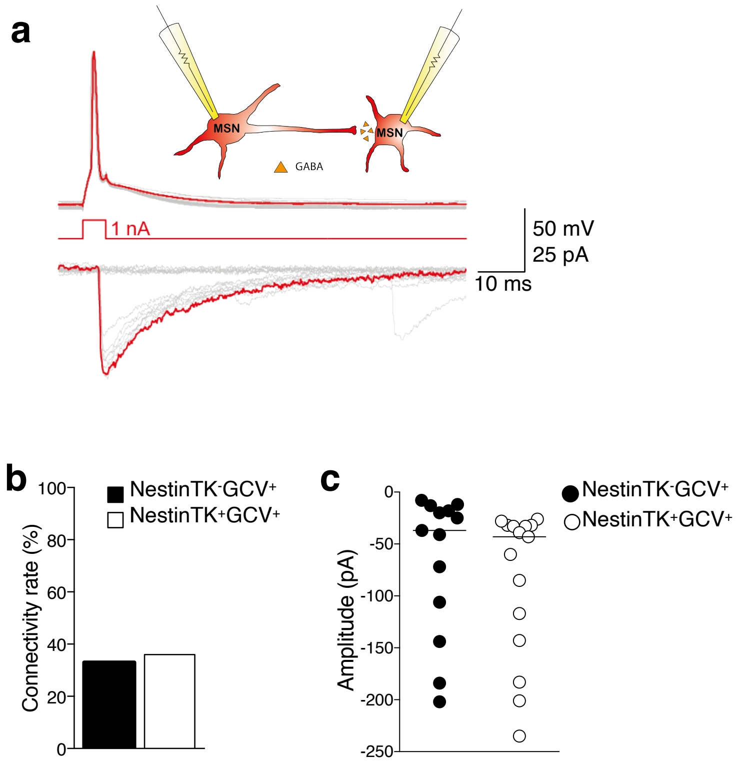
a
MSN
MSN
GABA
1 nA
50 mV
25 pA
10 ms
b
Connectivity rate (%)
NestinTK-GCV+
NestinTK+GCV+
c
Amplitude (pA)
NestinTK-GCV+
NestinTK+GCV+
d
Failure rate (%)
NestinTK-GCV+
NestinTK+GCV+

Figure S3

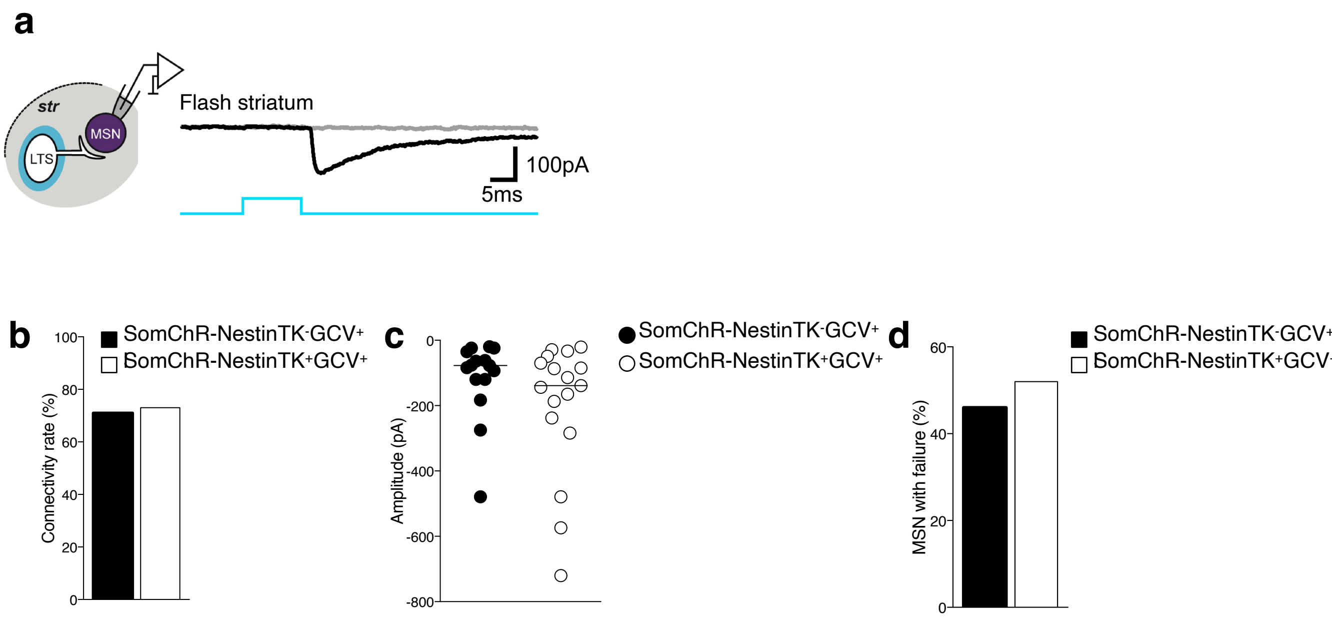
a
str
MSN
LTS
Flash striatum
100pA
5ms
b
SomChR-NestinTK-GCV+
SomChR-NestinTK+GCV+
Connectivity rate (%)
c
SomChR-NestinTK-GCV+
SomChR-NestinTK+GCV+
Amplitude (pA)
d
SomChR-NestinTK-GCV+
SomChR-NestinTK+GCV+
MSN with failure (%)

Figure S4

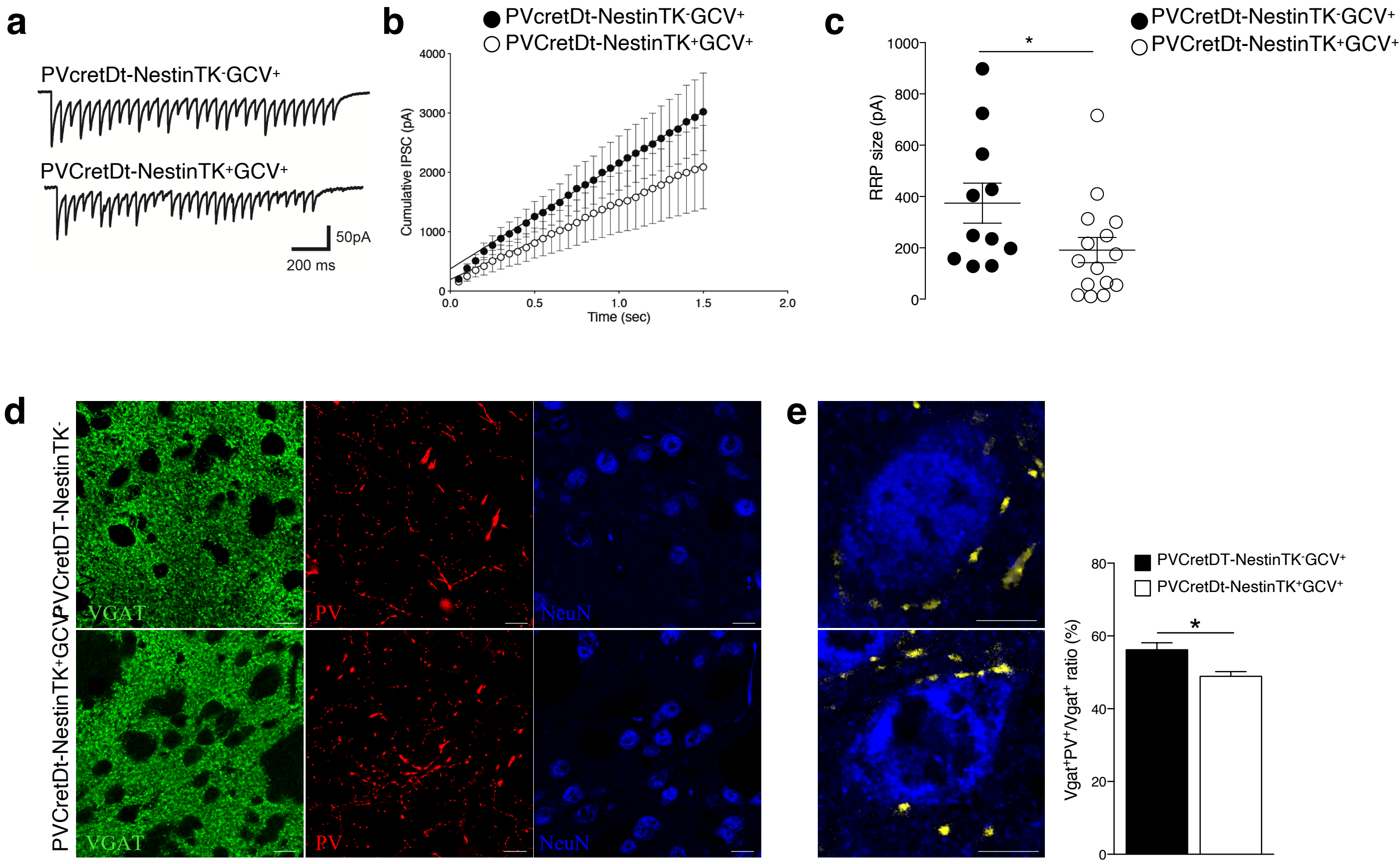

a
PVcretDt-NestinTK-GCV+
PVCretDt-NestinTK+GCV+
50pA
200 ms
b
PVcretDt-NestinTK-GCV+
PVCretDt-NestinTK+GCV+
Cumulative IPSC (pA)
Time (sec)
c
PVCretDt-NestinTK-GCV+
PVCretDt-NestinTK+GCV+
RRP size (pA)
*
d
PVCretDT-NestinTK-
PVCretDt-NestinTK+GCV+
VGAT
PV
NeuN
e
PVCretDT-NestinTK-GCV+
PVCretDt-NestinTK+GCV+
Vgat+PV+/Vgat+ ratio (%)
*


Figure S5

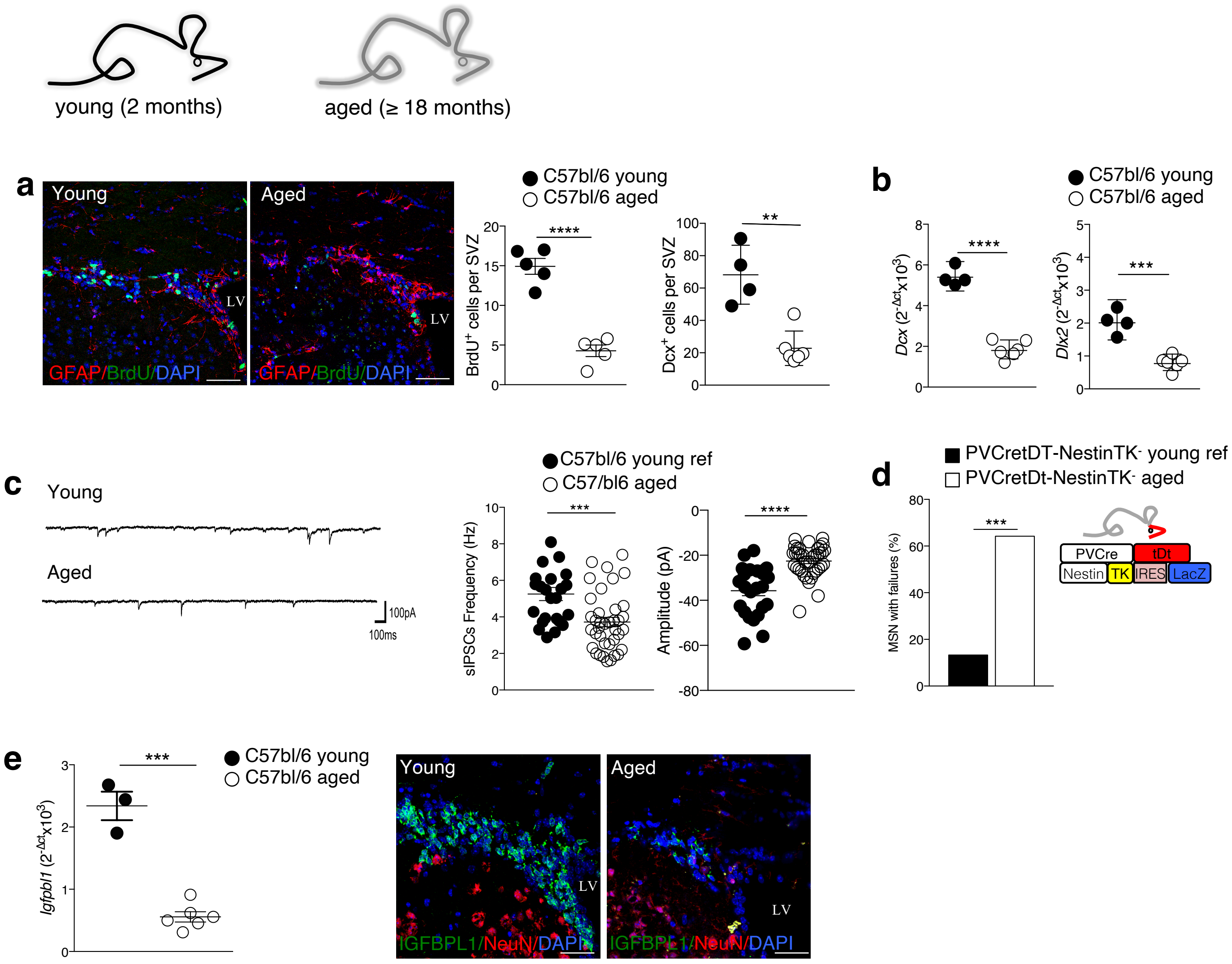
young (2 months)
aged (≥ 18 months)
a
Young
Aged
LV
GFAP/BrdU/DAPI
C57bl/6 young
C57bl/6 aged
BrdU+ cells per SVZ
Dcx+ cells per SVZ
b
Dcx (2-Δctx10³)
Dlx2 (2-Δctx10³)
c
Young
Aged
100pA
100ms
C57bl/6 young ref
C57/bl6 aged
sIPSCs Frequency (Hz)
Amplitude (pA)
d
PVCretDT-NestinTK- young ref
PVCretDt-NestinTK- aged
MSN with failures (%)
PVCre
tDt
Nestin
TK
IRES
LacZ
e
Igfpbl1 (2-Δctx10³)
IGFBPL1/NeuN/DAPI

Figure S6

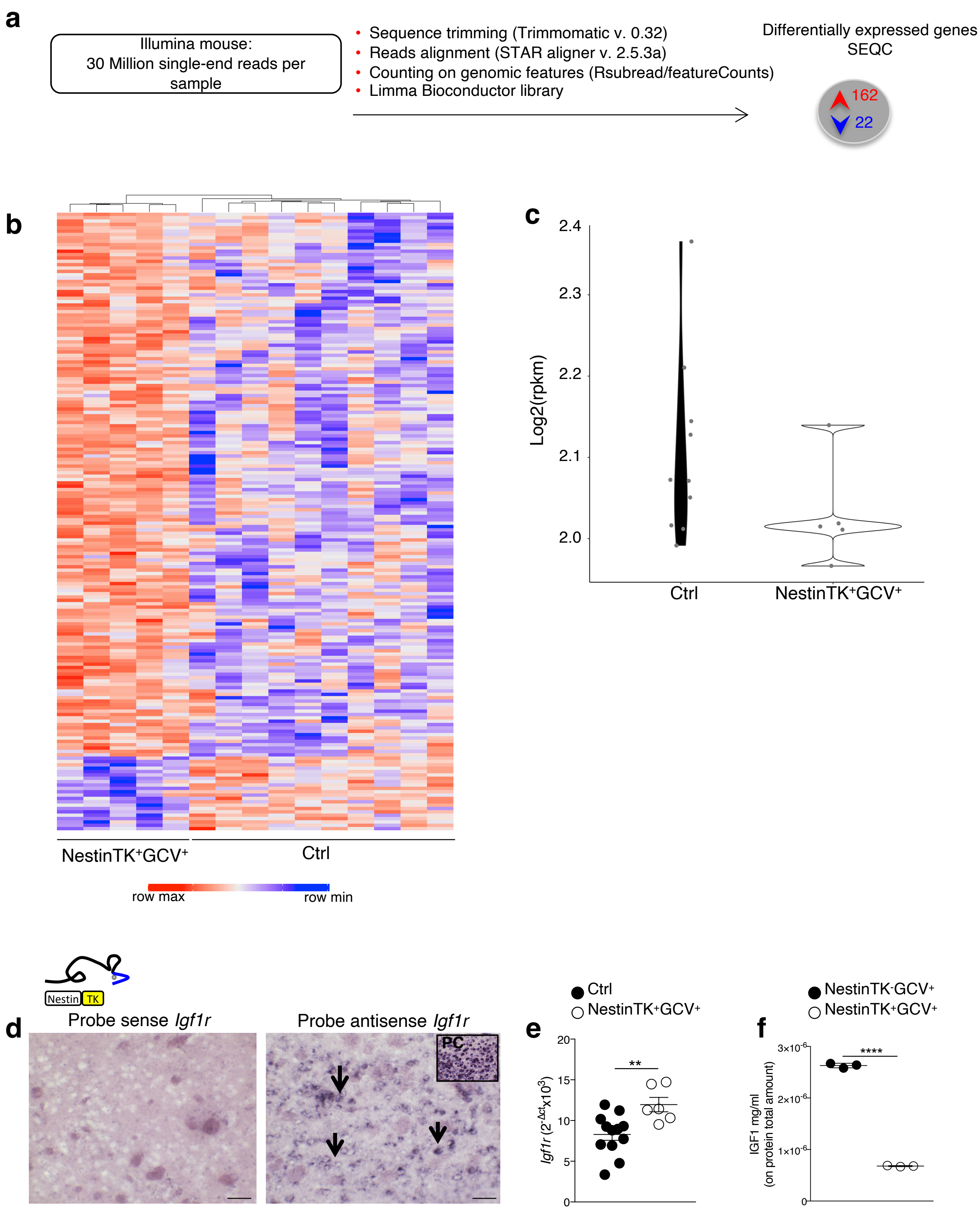
a
Illumina mouse:
30 Million single-end reads per sample
Sequence trimming (Trimmomatic v. 0.32)
Reads alignment (STAR aligner v. 2.5.3a)
Counting on genomic features (Rsubread/featureCounts)
Limma Bioconductor library
Differentially expressed genes
SEQC
162
22
b
NestinTK+GCV+
Ctrl
row max
row min
c
Log2(rpkm)
2.4
2.3
2.2
2.1
2.0
Ctrl
NestinTK+GCV+
Nestin
TK
d
Probe sense Igf1r
Probe antisense Igf1r
PC
e
Ctrl
NestinTK+GCV+
Igf1r (2-Δct x10³)
**
f
NestinTK-GCV+
NestinTK+GCV+
IGF1 mg/ml
(on protein total amount)
3x10-6
2x10-6
1x10-6
****

Figure S7
